# pH-Responsive Synthetic Cells for Programmable *in Situ* Protein Synthesis and Release

**DOI:** 10.1101/2025.11.16.688650

**Authors:** Sung-Won Hwang, Yudan Li, Alexander A. Green, Allen P. Liu

## Abstract

Lipid membrane-bound synthetic cells provide programmable, cell-like compartments for on-demand biomolecule production. However, most cell-free gene expression systems in synthetic cells are regulated by user-imposed cues rather than signals associated with environmental or physiological states. Here, we present an acidity-transducing synthetic cell that converts external pH changes into nucleic acid information to drive *in situ* protein synthesis. The system integrates gramicidin A proton channels for pH sensing, triplex-forming single-stranded DNA (ssDNA) that releases a trigger ssDNA upon acidification, and a toehold switch RNA that activates translation in response to the released trigger ssDNA. This work further reveals that tuning the annealing length between the pH-responsive and trigger ssDNAs controls the trigger-release pH, which is critical for enabling acid-triggered protein synthesis. The synthetic cells retain their pH-responsive activity when embedded in alginate hydrogels, creating acidity-responsive materials that synthesize proteins *in situ* rather than release preloaded cargo. Cell-penetrating peptide tagging further enables selective protein release and target-specific binding. This work establishes a molecular transduction strategy for programming synthetic cell materials to sense environmentally relevant acidity and generate functional biomolecules on demand.

**Table of Contents:** A synthetic cell is engineered to convert environmental acidification into nucleic acid signaling that activates protein synthesis. By coupling proton-channel sensing, pH-responsive DNA transduction, and toehold-regulated translation, the platform enables acid-triggered protein production within hydrogel-embedded synthetic cells. This strategy expands the repertoire of condition-associated cues that synthetic cells can interpret for responsive biomaterials, biosensing, and controlled delivery.

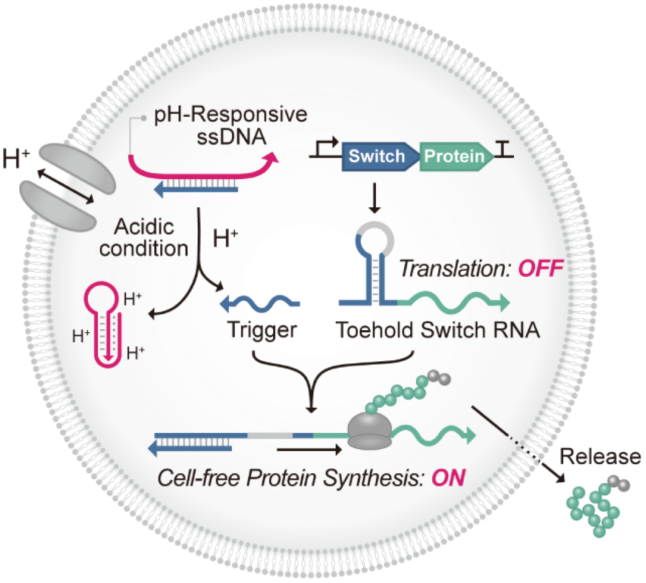

## 1. Introduction

Living cells possess sophisticated mechanisms for sensing and responding to environmental stimuli. However, engineering living cells to generate desired outputs in response to defined inputs remains challenging, as their native regulatory networks—of which we have only a partial understanding—can interfere with introduced synthetic components. Synthetic cells offer a bottom-up approach to overcome these limitations while avoiding risks associated with uncontrolled propagation.^[1]^ They are cell-sized systems assembled from well-defined, non-living components. Typically constructed as vesicle-based platforms with a phospholipid bilayer that mimics the membrane architecture of living cells, synthetic cells can incorporate membrane proteins for environmental sensing and drive protein synthesis through encapsulated cell-free transcription–translation machinery.^[2]^

Controlling gene expression in synthetic cells in response to external stimuli is essential for their use as biosensors and as on-demand production platforms that generate active biomolecules at the point of need. Yet most gene-control strategies in synthetic cells rely largely on user-imposed cues, including small-molecule inducers used in proof-of-concept riboswitches, such as theophylline,^[3–5]^ arabinose,^[5]^ and adenine,^[3]^ as well as externally applied physical inputs, including light^[6]^ and magnetic fields.^[7]^ By contrast, relatively few systems respond to signals intrinsically linked to biologically meaningful states, such as elevated temperature^[8]^ or environmentally and biologically relevant chemicals, including fluoride^[9]^ and histamine.^[10]^ Broadening this sensing repertoire—particularly toward signals that indicate conditions requiring timely intervention—is critical to enabling synthetic cells to autonomously respond in diverse, practically relevant settings.

Among such underexplored cues in synthetic cells, pH is particularly attractive because pH changes often mark critical transitions in biological systems. Under normal conditions, physiological pH is tightly regulated within a narrow range of 7.35–7.45, and gradual deviations toward acidosis (pH < 7.35) are hallmarks of several pathological conditions.^[11]^ Mildly acidic microenvironments are characteristic of tumors (pH 6.4–6.8),^[12,13]^ and inflamed surgical tissues (pH 6.0–6.8),^[14,15]^ as a result of hypoxia, elevated glycolytic metabolism, and lactic acid accumulation. Similarly, ischemia during myocardial infarction and stroke can rapidly lower local tissue pH to approximately 6.0–6.5.^[16]^ For targeting such acidic conditions, a variety of pH-responsive molecular delivery systems have been developed. Most of these systems, including pH-responsive hydrogels, rely on acid-sensitive chemical bonds or polymers that undergo bond cleavage or structural changes, thereby releasing preloaded cargo.^[17,18]^ However, such release-based approaches face inherent limitations, including passive leakage by diffusion and a lack of active, stimulus-dependent control over biomolecule production. The critical technological gap, therefore, lies in developing proactive systems capable of monitoring acidic stress and synthesizing functional biomolecules on demand directly at the site where they are needed.

Here, we report a pH-responsive synthetic cell capable of synthesizing and releasing proteins under acidic conditions. This system integrates three components (**Fig. 1a**): (i) the proton channel gramicidin A^[19]^ for pH sensing, (ii) a pH-responsive triplex-forming single-stranded DNA (ssDNA)^[20]^ that functions as a transducer, and (iii) a toehold switch RNA^[21]^ serving as an on–off regulator of protein synthesis. Gramicidin A is a bacterial peptide that forms a selective transmembrane ion channel in lipid membranes,^[19]^ enabling proton transport across vesicle membranes and thus allowing synthetic cells to sense external pH.^[22]^ The pH-responsive ssDNA consists of two intramolecular hairpins: a pH-insensitive hairpin stabilized by Watson–Crick base pairing and a pH-sensitive hairpin formed through parallel Hoogsteen interactions.^[20,23]^ At basic pH, a complementary ssDNA can hybridize with the exposed loop domain. Upon acidification, the Hoogsteen interactions are strengthened, leading to the formation of an intramolecular triplex structure that displaces and releases the complementary ssDNA.^[24,25]^ The toehold switch is a *de novo*-designed riboregulator that inhibits protein expression by forming a hairpin structure around the ribosome binding site (RBS), thereby preventing translation initiation. It switches from OFF to ON state and allows protein translation when a cognate trigger binds and unfolds the hairpin, exposing the RBS.^[21]^ Unlike riboregulators inspired by natural systems, toehold switches impose few sequence restrictions on the trigger strand, making them highly adaptable to diverse DNA-based inputs. In our system, the ssDNA released from the pH-responsive ssDNA under acidic conditions serves as the trigger that activates the toehold switch, enabling acid-triggered protein synthesis.

**Fig. 1.**
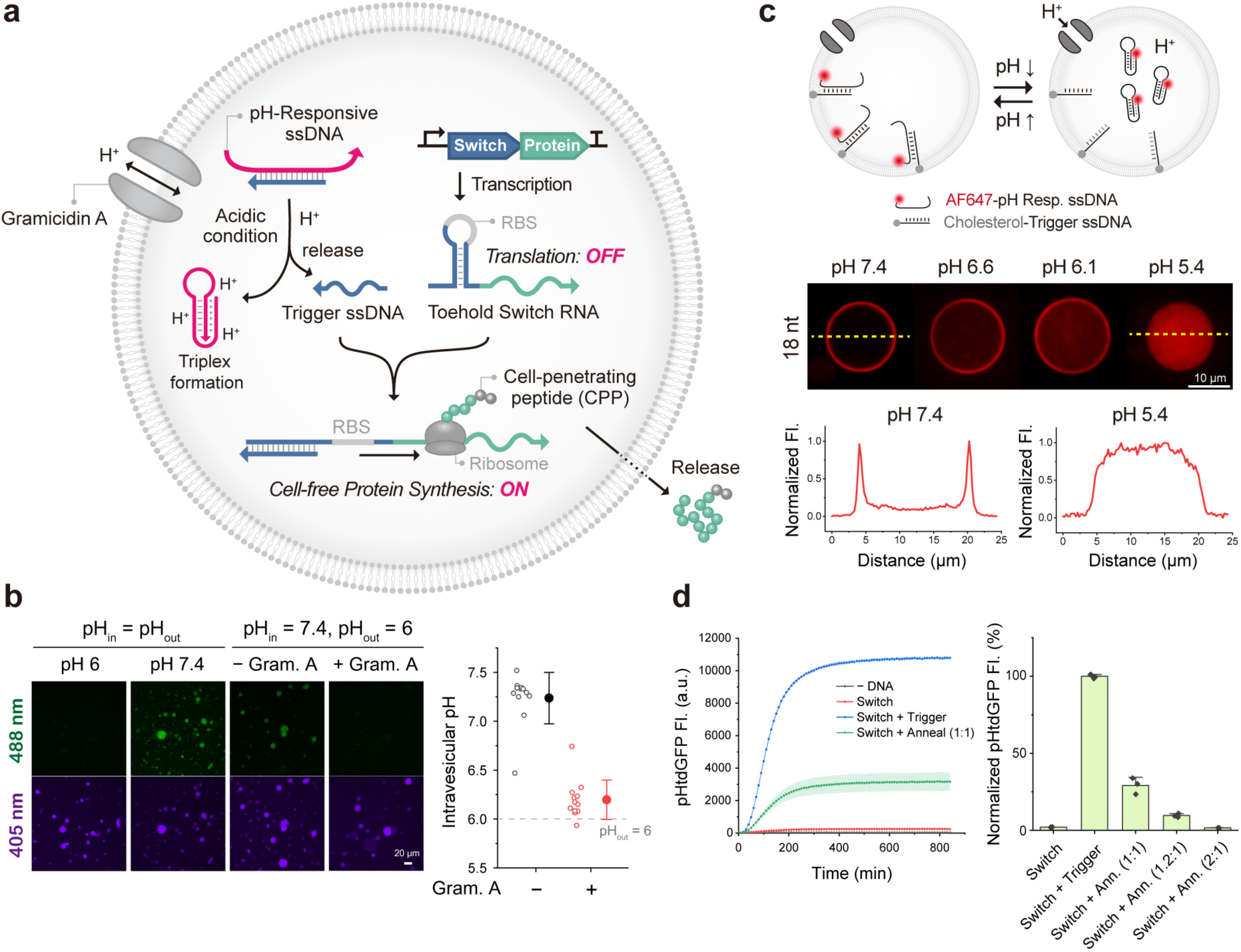
Engineering a pH-responsive synthetic cell. **a**, Schematic of acid-responsive cell-free protein expression inside a synthetic cell. At pH 7.4, the toehold switch inhibits protein expression by forming a hairpin structure around the ribosome binding site (RBS). At acidic pH, protons diffuse into the synthetic cell through gramicidin A channels, and the pH-responsive single-stranded DNA (ssDNA) forms a triplex structure, releasing a complementary trigger ssDNA. The released trigger strand binds to the toehold switch, unwinds the hairpin, and enables protein synthesis. **b**, Gramicidin A is required for intravesicular acidification. Intravesicular pH was measured using encapsulated HPTS (pyranine) in synthetic cells initially at pH_in_ = 7.4 exposed to pH_out_ = 6.0 for 14 h, with or without gramicidin A (calibration in Supplementary Fig. 2). Without gramicidin A, intravesicular pH remained near neutral (7.2 ± 0.3); with gramicidin A, it dropped to 6.2 ± 0.2, close to pH_out_, demonstrating that gramicidin A is required for efficient coupling of external and internal pH. Open circles, individual GUVs; filled circles, mean ± s.d. (*n* = 12 vesicles per condition). **c**, Trigger ssDNA release inside a synthetic cell under acidic pH. To distinguish between the hybridized or detached states of the pH-responsive DNA and trigger DNA, they were labeled with Alexa Fluor 647 (AF647) and cholesterol, respectively, and encapsulated inside synthetic cells with gramicidin A channels. Since the trigger ssDNA remains at the vesicle membrane via cholesterol, AF647 signal observed on the membrane under neutral pH indicates the hybridized state, whereas AF647 fluorescence in the lumen under acidic pH indicates the detached state, as illustrated in the representative fluorescence images and associated line scan analysis. **d**, Toehold switch– mediated cell-free protein expression. (Left) Toehold switch without or with trigger ssDNA present OFF/ON expression of pHtdGFP. The Anneal (Ann.) condition refers to hybridized pH-responsive ssDNA + trigger ssDNA. (Right) Trigger ssDNA supplied as a hybridized complex with pH-responsive ssDNA (1:1 ratio) results in leaky expression, which is reduced by increasing the ratio of the pH-responsive strand to 2:1. Fluorescence is quantified at 14 h and normalized to the Switch + Trigger condition. Error bars are mean ± s.d. (*n* = 3 independent cell-free reactions).

As a demonstration, we synthesized NanoLuc (NLuc) luciferase, and a pH-stable tandem dimeric GFP (pHtdGFP).^[26]^ Unlike conventional GFP, pHtdGFP contains two point mutations (N149Y and Q204H) that have been proposed to restrict proton access to the chromophore, thereby maintaining its fluorescence under acidic conditions.^[26]^ We also identified the annealing length between the pH-responsive and trigger ssDNAs as a molecular design parameter that tunes the pH threshold, thereby matching the DNA transduction module to the operational pH range of cell-free expression. Finally, we embedded pH-responsive synthetic cells in alginate hydrogels to generate acidity-gated, protein-producing biomaterials. Incorporating a cell-penetrating peptide (CPP) strategy further demonstrated the export of synthesized proteins and target binding. Together, these results establish a platform for integrating biologically relevant acidity into programmable synthetic cells, with potential applications in biosensing, responsive biomaterials, and controlled delivery in therapeutic contexts.

## 2. Results and Discussion

### 2.1. Synthetic cells transduce external acidification into trigger ssDNA release

We first examined whether gramicidin A was required to transmit external acidification across the synthetic cell membrane. To directly monitor intravesicular pH, we encapsulated the pH-sensitive fluorescent dye 8-hydroxypyrene-1,3,6-trisulfonic acid trisodium salt (HPTS) in vesicles. HPTS exhibits pH-dependent protonation of its 8-hydroxyl group (pKa = 7.22), with emission at 511 nm increasing with pH upon excitation near 460 nm and decreasing upon excitation near 406 nm.^[22,27]^ HPTS (100 μM) was encapsulated in vesicles prepared at pH values ranging from 5 to 9 (using Tris-, HEPES-, and MES- solutions) using an inverted emulsion method^[28]^ (**Supplementary Fig. 1**) and imaged under confocal microscopy using 488- and 405-nm excitation. Fluorescence intensities were analyzed to generate a calibration curve (**Supplementary Fig. 2**).

Vesicles prepared at pH 7.4 were exposed to an external solution at pH 6.0 in the presence or absence of 10 μg/mL gramicidin A. After 14 h, vesicles containing gramicidin A showed substantially decreased 488-nm HPTS fluorescence, comparable to that of vesicles equilibrated at pH 6.0 (**Fig. 1b**). pH values estimated from the calibration curve further showed that most gramicidin A-containing vesicles had acidified close to pH 6.0. In contrast, vesicles without gramicidin A retained 488-nm fluorescence, indicating that external acidification was not efficiently transmitted into the vesicle lumen. These results establish that gramicidin A enables equilibration of intravesicular pH with the acidic external environment.

Having established gramicidin A-mediated intravesicular acidification, we next constructed the nucleic acid transduction module using a pH-responsive ssDNA and a complementary trigger ssDNA, based on previous studies^[24,25]^ with slight modifications. The pH-responsive ssDNA contains a hairpin loop and a pH-sensitive stem domain (**Supplementary Fig. 3** and **Supplementary Table S1**). The trigger strand was designed to be partially (18 nt) complementary to the loop domain, allowing it to hybridize with the pH-responsive ssDNA at basic pH. Under acidic conditions, the stem domain forms a triplex that destabilizes this hybridization, leading to trigger strand release.

To visualize this process in synthetic cells, the 5’ end of the pH-responsive ssDNA and the 3’ end of the trigger ssDNA were labeled with Alexa Fluor 647 (AF647) and cholesterol, respectively, and the two strands were hybridized (see Methods). The annealed constructs were encapsulated in vesicles via the inverted emulsion method. Because the cholesterol-labeled trigger strand remains in the vesicle membrane through the cholesterol anchor,^[29]^ the AF647 signal of the pH-responsive strand could be identified as being annealed to the trigger if it is present on the membrane or detached if it is present in the lumen^[24]^ (**Fig. 1c**).

Synthetic cells initially at pH 7.4 were exposed to various pH conditions ranging from pH 7.8 to 5.4 in the presence of 10 μg/mL gramicidin A. At pH 5.4, the AF647 signal shifted from the membrane to the lumen, indicating that pH-responsive ssDNA with an 18-nt annealing length is detached from the trigger strand (**Fig. 1c**). This confirms successful trigger release in response to acidification and demonstrates that synthetic cells can use gramicidin A channels to sense external pH through proton influx.

### 2.2. Toehold switches enable cell-free translational regulation by trigger ssDNA

After confirming successful trigger strand release from the pH-responsive ssDNA upon acidification, we next designed the toehold switch, which is activated by the trigger ssDNA. To prevent activation at neutral pH, the exposed toehold region complementary to the trigger ssDNA was made shorter than the 18-nt complementary region of the pH-responsive ssDNA, giving the trigger ssDNA a lower affinity for the toehold switch than for the pH-responsive ssDNA under neutral conditions. 25 toehold switch candidates with varying stem and toehold lengths were generated using a NUPACK-based sequence selection algorithm.^[21]^ Bulk cell-free expression of pHtdGFP using the PUREfrex was tested to screen these candidates based on the ON/OFF ratios in the presence or absence of the trigger ssDNA. A switch with a total trigger-binding length of 26 nucleotides was selected, consisting of a 9-nt toehold region and a hairpin with 6-bp and 8-bp stems flanking a 3-nt bulge (**Supplementary Fig. 3a–b**). This switch exhibited negligible expression without the trigger (∼2% of that observed with trigger), but robust activation upon trigger addition (**Fig. 1d**, left). However, when the trigger ssDNA pre-annealed with the pH-responsive ssDNA (at a 1:1 ratio) was added, partial expression (∼29.1%) occurred. To reduce this leaky activation, the ratio of pH-responsive ssDNA to trigger ssDNA was adjusted. Increasing the ratio to 2:1 minimized leakage (∼1.7%), comparable to the no-trigger control (**Fig. 1d**, right).

### 2.3. Annealing length controls acid-responsive protein expression in synthetic cells

We next assembled the toehold switch plasmid with the annealed pH-responsive and trigger ssDNAs and tested protein expression in synthetic cells, but acid-responsive protein synthesis was not observed. To identify the cause, we first determined the functional pH range of toehold-mediated expression using synthetic cells encapsulating an NLuc-encoding toehold switch plasmid and the trigger ssDNA, but without the pH-responsive ssDNA, allowing the trigger to constitutively activate translation. The vesicles containing gramicidin A were incubated at 37 °C for 20 h at pH 7.4, 7.0, 6.5, 6.0, and 5.5 to ensure a stable endpoint for protein-expression measurements (time-lapse pHtdGFP fluorescence approached a plateau by approximately 8 h; **Supplementary Fig. 4**). Samples were then adjusted to the same final pH of 7.3 to minimize pH-dependent differences in NLuc activity, and luminescence was measured after addition of a membrane-permeable^[30,31]^ NLuc substrate furimazine (see Methods and **Supplementary Fig. 5a**). Luminescence peaked at pH 7.0 and decreased with increasing acidity, with almost no expression at pH 5.5 (**Fig. 2a**, left), likely due to reduced activity of transcription–translation enzymes. In contrast, the annealed pH-responsive and trigger ssDNAs exhibited a transition midpoint (pH1/2) of 6.2, with noticeable trigger release (luminal signal) observed at pH 5.4 (**Fig. 1c**), indicating that the trigger release occurs in a pH range where protein synthesis is severely limited.

**Fig. 2.**
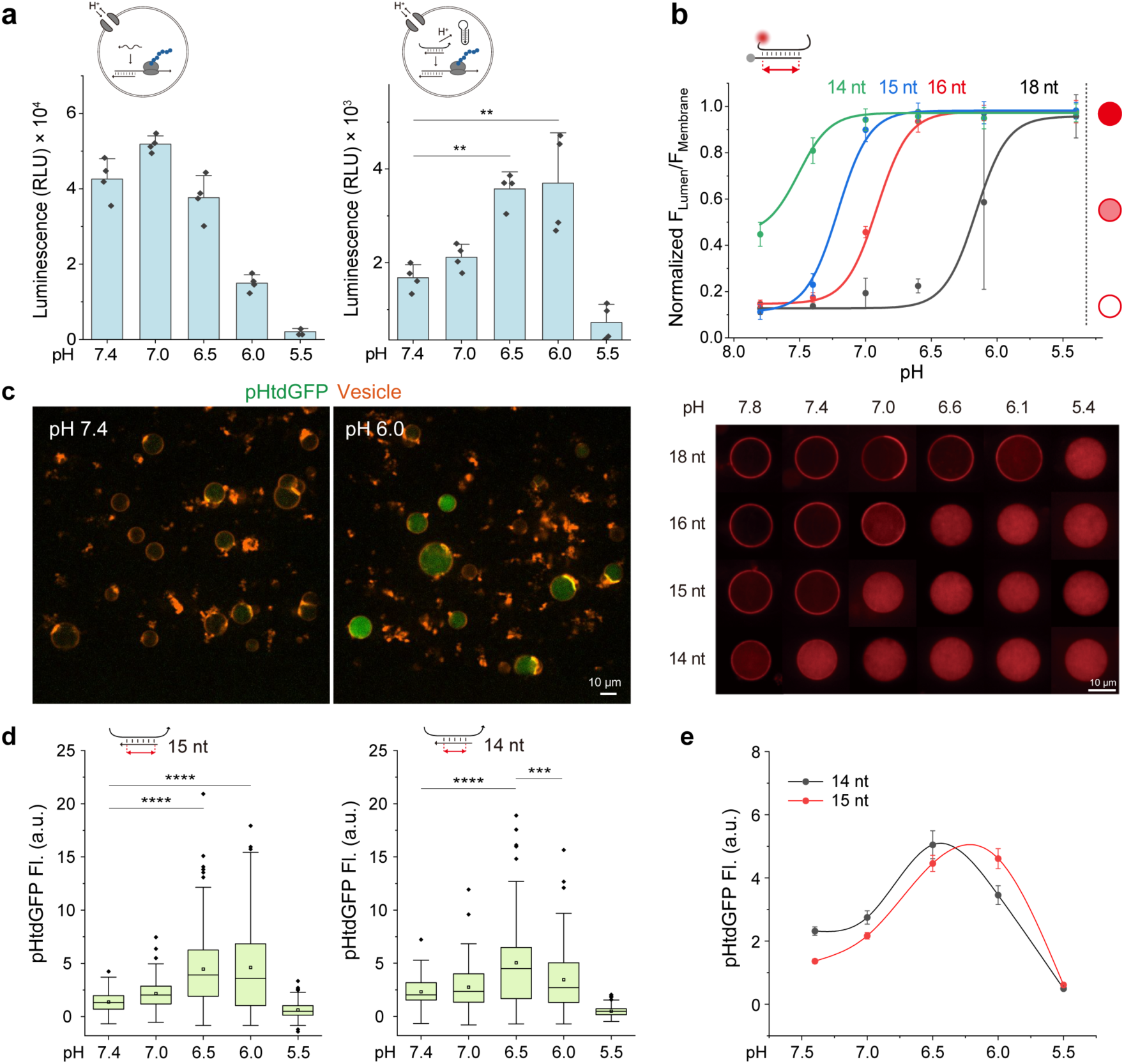
Tuning the annealing length of pH-responsive and trigger ssDNAs enables acid-triggered protein synthesis in synthetic cells. **a**, Luminescence of NanoLuc (NLuc) expressed inside synthetic cells under different pH conditions (7.4, 7.0, 6.5, 6.0, and 5.5). (Left) Without pH-responsive ssDNA, synthetic cells containing trigger ssDNA and toehold switch-NLuc DNA show the highest expression at neutral pH. (Right) With 15-nt annealing length pH-responsive ssDNA, maximal luminescence occurs at pH 6.0, indicating acid-responsive protein synthesis. Statistical significance was determined by ANOVA: ** *p* < 0.01 (*n* = 4 independent experiments). Error bars are mean ± s.d. **b**, (Top) Shorter annealing length increases the pH transition midpoint (pH_1/2_) at which fluorescence shifts from the vesicle membrane to the lumen, indicating the transition from the hybridized to the detached state. Error bars are mean ± s.d. (*n* = 6–10 vesicles per condition). (Bottom) Representative fluorescence images of AF647 pH-responsive ssDNA with trigger ssDNA from 14-nt to 18-nt annealing lengths, incubated between pH 7.8 and 5.4. **c**, Confocal microscopy images of pHtdGFP expression in synthetic cells at pH 7.4 and pH 6.0 with 15-nt annealing length of pH-responsive ssDNA and trigger ssDNA. **d**, Comparison of pHtdGFP fluorescence in synthetic cells with 15-nt (left) or 14-nt (right) annealing length across different pH values. Statistical significance was determined by ANOVA: *** *p* < 0.001; **** *p* < 0.0001 (*n* = 98–182 individual vesicles). Box plot shows the 25–75th percentile range; the central line indicates the median, squares mark the mean, whiskers extend to data points within 1.5 times the interquartile range, and outliers are depicted individually. **e**, Trendline of average fluorescence values from **d** shows that decreasing the annealing length from 15-nt to 14-nt shifts the expression peak toward milder acidic pH. Error bars indicate standard error of the mean.

Therefore, we sought to shift the trigger-release transition midpoint (pH1/2) to a slightly higher, yet still acidic, pH range, and tested several strategies—including adjusting TAT% in the pH-responsive stem,^[23]^ varying ssDNA and plasmid concentrations,^[32]^ and introducing mismatches—but these mainly caused leaky expression under neutral conditions. Ultimately, we found that tuning the annealing length between the pH-responsive and trigger ssDNAs effectively shifted the transition pH. Reducing the annealing length from 18 to 16, 15, and 14 nucleotides gradually increased the pH1/2, enabling trigger release under milder acidity (**Fig. 2b**; **Supplementary Fig. 6a-c; Supplementary Table S2**). Within the 14–18 nt range examined, quantitative analysis yielded a slope of −0.303 ± 0.012 pH units per nucleotide (R^2^ = 0.997; **Supplementary Fig. 6d**; see Methods for line scan analysis and **Supplementary Fig. 7**), corresponding to an approximately 0.30-pH-unit increase in the transition midpoint for each nucleotide shortened. As expected, gramicidin A-mediated pH equilibration was necessary for the detachment of pH-responsive ssDNA with a shorter annealing length under acidic conditions (**Supplementary Fig. 8**).

When the reduced 15-nt annealing construct was tested with the toehold switch for NLuc expression in synthetic cells (**Supplementary Fig. 3c–d**), the highest expression was observed at pH 6.0 (**Fig. 2a**, right), confirming successful acid-triggered gene activation. Normalization to the corresponding trigger control at each pH further showed increased relative activation of the pH-responsive system under more acidic conditions (**Supplementary Fig. 5b**). To examine expression at the single-vesicle level, pHtdGFP was expressed. Synthetic cells encapsulating the toehold–pHtdGFP plasmid and the 15-nt construct showed the strongest signal at pH 6.0, which was 3.4-fold higher than at pH 7.4 (**Fig. 2c; 2d**, left; **Supplementary Fig. 9**). The 14-nt construct also showed acid-responsive behavior, with 2.2- fold higher maximal expression at pH 6.5 (**Fig. 2d–e**), reflecting its higher transition midpoint compared with the 15-nt construct. This demonstrates that tuning the annealing length between the pH-responsive and trigger ssDNAs enables control of the target pH for protein expression. However, shorter annealing lengths increased leaky expression in bulk, which was mitigated by increasing the pH-responsive:trigger ratio from 2:1 to 3:1 (**Supplementary Fig. 10**). We selected the 15-nt construct at a 3:1 ratio for subsequent experiments as it showed the largest fold change relative to the pH 7.4 control. Nevertheless, the lower pH limit of the current system remains constrained by the pH tolerance of the cell-free expression machinery, which exhibits substantially reduced activity below approximately pH 6.0. Extending this approach toward lower pH values would therefore require future engineering or reformulation of the cell-free expression system, for example, by improving low-pH-sensitive components through protein engineering^[33]^ or replacing them with functionally equivalent homologs from acid-tolerant organisms.^[34]^

### 2.4. Embedding pH-responsive synthetic cells creates acidity-gated protein-producing hydrogels

Having established acidity-to-protein-synthesis transduction in vesicles, we next asked whether these synthetic cells could endow a bulk soft material with acidity-gated biomolecule production. Acid sensing and trigger release occur rapidly, with ∼73.5% strand detachment within 10 min (**Supplementary Fig. 11**), whereas cell-free expression proceeds more slowly, requiring approximately 2 h to reach half of the maximum yield (**Fig. 1d**). Immobilizing synthetic cells in a hydrogel is therefore advantageous because it spatially retains the vesicles during the protein expression while allowing diffusion-based access to protons and small-molecule substrates. More importantly, hydrogels provide a three-dimensional material format that may facilitate the future development of localized, responsive biomaterials for biomedical applications. Embedding synthetic cells also offers a modular approach to creating functional hydrogels. We previously demonstrated that incorporating stimuli-responsive materials into hydrogels confers responsiveness without altering the hydrogel’s chemical structures.^[35]^ Similarly, pH-responsive synthetic cells can serve as embedded reaction modules, transforming an otherwise passive hydrogel into a protein-producing material. Although many acid-responsive hydrogels that release preloaded payloads have been reported,^[17,36]^ *in situ* acid-triggered protein synthesis has not yet been demonstrated. The synthetic cells introduced here thus create a new class of biomaterial capable of synthesizing proteins on demand in response to acidity.

Before evaluating protein expression, we characterized vesicle loading and stability within the hydrogel. Vesicles encapsulating 2 µM Cy5 dye were mixed with alginate to a final concentration of 0.8% w/v and crosslinked by calcium ions to form vesicle-containing hydrogels^[35]^ (see Methods). Confocal Z-stack analysis showed an embedded vesicle density of approximately 1.22 × 10^4^ vesicles/µL of hydrogel, and a vesicle volume fraction of 0.23 ± 0.04%, and their three-dimensional distribution is shown in **Supplementary Video 1**. The embedded vesicles were further monitored at room temperature for 8 days. Approximately 71% and 83% of the tracked vesicles retained Cy5 fluorescence at pH 7.4 and pH 6.5, respectively (**Supplementary Fig. 12**). Notably, at pH 6.5, only ∼40% of vesicles in solution retained Cy5 fluorescence by day 8, compared with ∼83% of hydrogel-embedded vesicles, suggesting that hydrogel embedding may improve vesicle stability under acidic conditions.

We next tested whether synthetic cells retained acid-responsive protein expression after hydrogel embedding. Synthetic cells containing acid-responsive protein expression components (toehold switch–pHtdGFP plasmid and hybridized pH-responsive and trigger ssDNAs, with 15-nt annealing length), together with gramicidin A, were mixed with alginate and crosslinked (**Fig. 3a–b**). Two different conditions (pH 7.4 and pH 6.5) of hydrogels containing synthetic cells were incubated at 37 °C for 20 h and imaged by confocal microscopy. As expected, embedded pH-responsive synthetic cells exhibited negligible pHtdGFP fluorescence at pH 7.4 but strong signals at pH 6.5 (**Fig. 3c–d**; **Supplementary Fig. 13a**), although the fluorescence intensity was significantly lower in the hydrogel than in solution (**Supplementary Fig. 13b**). These results demonstrate that synthetic cells retain pH responsiveness and protein synthesis capability within a hydrogel, although the overall expression level is reduced, highlighting their capacity to create pH-responsive biomaterials.

**Fig. 3.**
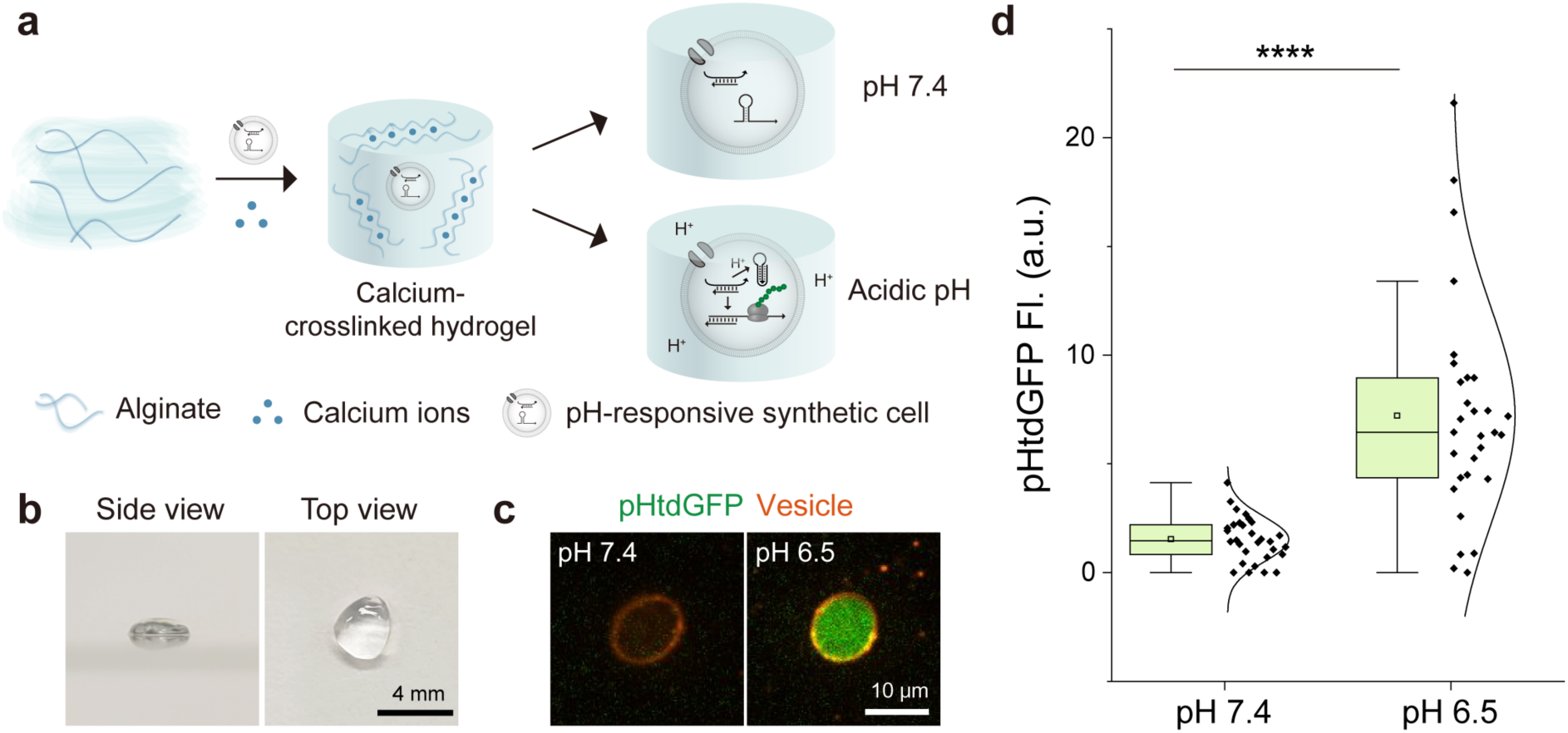
Embedding pH-responsive synthetic cells in alginate hydrogels creates acidity-gated protein-producing biomaterials. **a**, Schematic of hydrogel formation. 1.5% w/v alginate macromer solution was mixed with pH-responsive synthetic cells, and 50 mM calcium ions were added to crosslink the hydrogel. Protein expression from the embedded synthetic cells is triggered by acidic conditions. **b**, Photographs of hydrogels containing pH-responsive synthetic cells. **c**, Representative confocal images of embedded synthetic cells under different pH conditions. **d**, Normalized fluorescence intensity of synthetic cells in hydrogels under different pH conditions. Expression of pHtdGFP is significantly higher at pH 6.5. Statistical significance determined by two-tailed Student’s *t*-test: **** *p* = 9.28×10^-7^ (*n* = 30 vesicles per condition). Box plot shows the 25–75th percentile range; the central line indicates the median, squares mark mean values, whiskers extend to data points within 1.5 times the interquartile range.

Although applications in biological tissues were not evaluated in the present study, a three-dimensional hydrogel material format may support the future development of localized, responsive biomaterials. Further studies will be required to evaluate their biocompatibility, stability, localization, and functional protein production under physiologically relevant conditions. Protein diffusion also remains an important consideration. Combining this approach with hydrogel degradation—for example, by co-expressing degradative enzymes^[37,38]^ and tuning network mesh size or density—could enhance biomolecule release from the hydrogel.^[39]^

### 2.5. Protein release from synthetic cells via cell-penetrating peptides

In synthetic cell systems, cargo release is commonly achieved by pore formation, such as α-hemolysin,^[5,7,8,10,40]^ or by membrane degradation by enzymes such as phospholipase A.^[10,40]^ However, these methods are limited either by the small pore size, which restricts cargo size, or by uncontrolled release caused by membrane disruption, precluding subsequent processes. To avoid these limitations, we employed the cell-penetrating peptide (CPP) penetratin, a 16- amino-acid sequence (RQIKIWFQNRRMKWKK) derived from the third α-helix of the homeodomain of Antennapedia, a homeotic transcription factor in *Drosophila melanogaster*.^[41]^ As a cationic CPP, penetratin interacts with negatively charged membrane components, including anionic lipids,^[42]^ and facilitates passive, concentration gradient-driven translocation of cargo across the bilayer.^[43]^ Unlike pore-forming peptides, penetratin translocates protein cargos as large as 100 kDa across membranes without inducing leakage.^[41,44]^ Recent work by Heili *et al.* demonstrated that penetratin-tagged proteins can be cell-free expressed and released from vesicle-based synthetic cells,^[43]^ highlighting their suitability for controlled protein transport.

Combining CPPs with targeting elements could enable acid-triggered, target-specific delivery of synthesized proteins. As a proof of concept, we tested whether proteins synthesized inside synthetic cells could be released and subsequently captured by external targets. We first constructed a toehold switch-controlled super-folder GFP (sfGFP) construct containing penetratin CPP at the N-terminus and a His-tag at the C-terminus (toehold switch- CPP-sfGFP-His) and tested its cell-free expression in bulk in the presence of the trigger strand. Compared with the non-CPP-tagged construct (toehold switch-sfGFP-His), the CPP-tagged construct showed robust sfGFP fluorescence **(Supplementary Fig. 14a)**. We then tested expression in synthetic cells and introduced nickel-nitriloacetic acid (Ni-NTA)-coated magnetic beads to capture released proteins via His-Ni-NTA binding **(Fig. 4a)**. Synthetic cells containing the toehold switch-CPP-sfGFP-His DNA, trigger ssDNA, and gramicidin A were prepared with the anionic lipid POPG (palmitoyl-2-oleoyl-sn-glycero-3-phospho-glycerol) incorporated in the membrane to facilitate interaction with the cationic penetratin peptide.^[43,45]^ After incubation at 37 °C for 20 h, vesicles expressing CPP-sfGFP-His exhibited membrane-localized fluorescence, while sfGFP-His alone remained luminal **(Supplementary Fig. 14b),** indicating penetratin-mediated membrane interaction. Upon bead addition, sfGFP fluorescence appeared on bead surfaces **(Fig. 4b–c; Supplementary Fig. 15.** Method for quantification, **Supplementary Fig. 16)**, confirming CPP-mediated protein release and binding to external targets. In contrast, no signal was detected from sfGFP-His-expressing controls lacking the CPP tag.

**Fig. 4.**
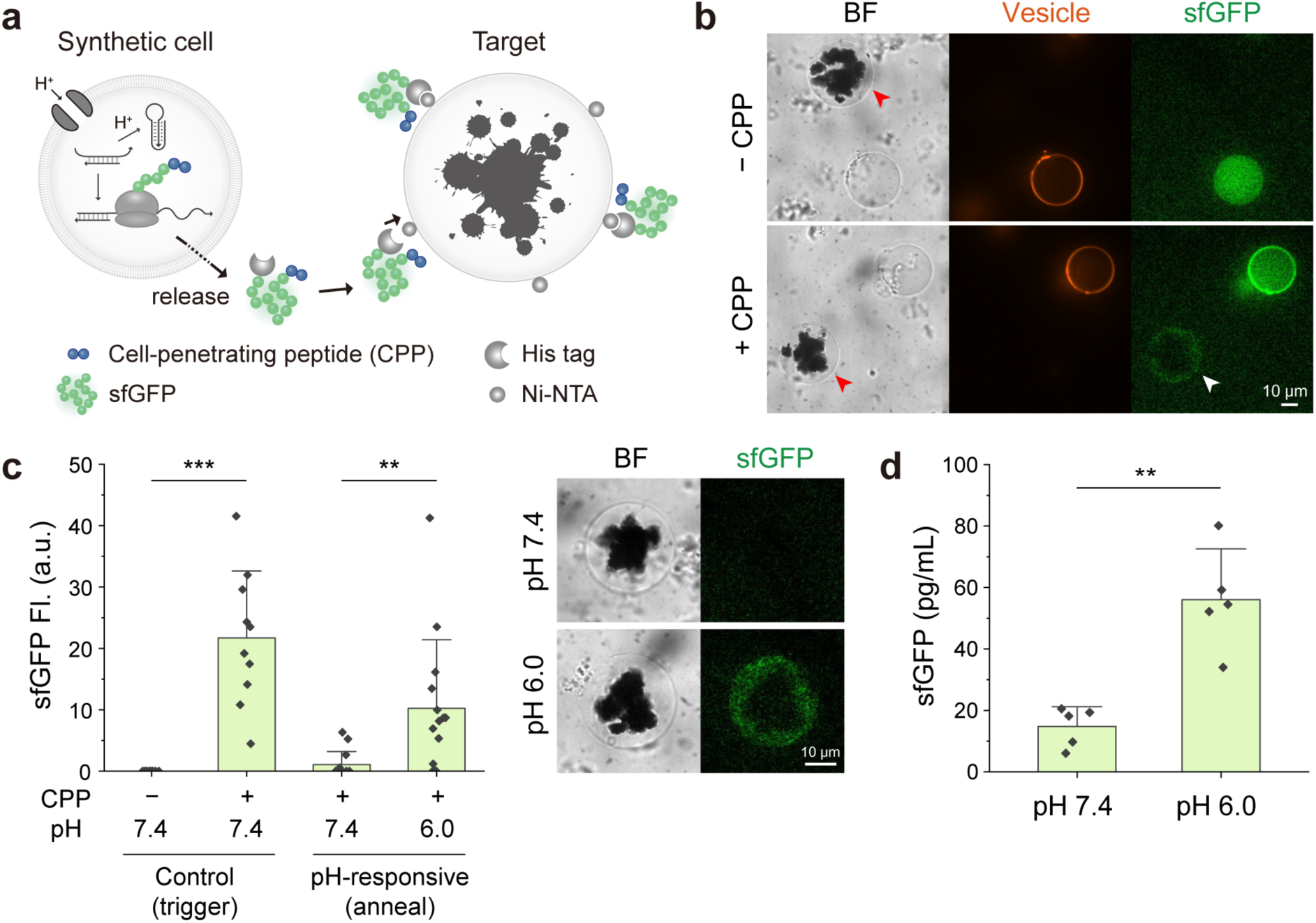
A cell-penetrating peptide (CPP) enables selective export and target-specific binding of proteins synthesized in synthetic cells. **a**, Schematic illustration of target-specific protein delivery from pH-responsive synthetic cells. Proteins expressed with both a CPP-tag and a His-tag can traverse the vesicle membrane and bind to Ni-NTA-functionalized targets via His-Ni-NTA interactions. **b**, Confocal fluorescence images of Ni-NTA-coated magnetic beads added to synthetic cells expressing sfGFP-His (−CPP) or CPP-sfGFP-His (+CPP). sfGFP fluorescence was observed only on bead surfaces incubated with CPP-sfGFP-His-expressing synthetic cells (white arrow). Ni-NTA-coated magnetic beads are indicated by red arrowheads in brightfield (BF) images. Synthetic cells (vesicles) are labeled with rhodamine–PE (orange). Membrane localization of sfGFP in vesicles indicates interactions between the cationic CPP and negatively charged POPG-containing lipid membranes. **c**, (Left) Quantification of sfGFP fluorescence on bead surfaces. Statistical significance was determined by a two-tailed Student’s *t*-test: *** *p* = 1.4×10^-4^; ** *p* = 9.2×10^-3^ (*n* = 10 beads for trigger conditions, 14 beads for anneal conditions). Error bars are mean ± s.d.. Anneal condition refers to hybridized pH-responsive ssDNA + trigger ssDNA. (Right) Representative confocal images of Ni-NTA-coated magnetic beads added to pH-responsive synthetic cells expressing CPP-sfGFP-His in pH 7.4 and pH 6.0. **d**, Quantification of released sfGFP by enzyme-linked immunosorbent assay (ELISA). Statistical significance was determined by a two-tailed Student’s *t*-test: ** *p* = 3.5×10^-3^ (*n* = 5 independent samples). Error bars are mean ± s.d..

We then applied this strategy to the acid-responsive system using the annealed construct (pH-responsive ssDNA hybridized with the trigger ssDNA with 15-nt annealing length) instead of the free trigger strand. As expected, sfGFP fluorescence on Ni–NTA bead surfaces was higher at pH 6.0 than at pH 7.4 (**Fig. 4c**). To quantitatively assess protein release, the sfGFP concentration in the external solution was measured by an enzyme-linked immunosorbent assay (ELISA) using the collected supernatant. Under pH 6.0 conditions at 20 h, released sfGFP was measured to be 56 pg/mL, which was approximately 3.8-fold higher than at pH 7.4 **(Fig. 4d)**. In a separate experiment, quantification of the released and vesicle-retained sfGFP fractions yielded a protein release efficiency of approximately 15.7% (**Supplementary Fig. 17**). These results confirm that CPP-assisted transport enables controlled, pH-dependent protein release from synthetic cells. Although not demonstrated in the present study, combining this CPP-mediated protein release module with hydrogel-embedded synthetic cells could potentially enable acidity-triggered protein production and subsequent release from the material.

## 3. Conclusions

Unlike living cells, synthetic cells are fully controllable, offering broad functional tunability.^[1]^ Here, we integrated three molecular modules—a proton channel, a pH-responsive triplex-forming ssDNA, and a toehold switch RNA—to construct a synthetic cell capable of acid-triggered protein synthesis and release. Whereas conventional acid-responsive systems are limited to releasing preloaded payloads, our platform enables on-demand synthesis of functional biomolecules in response to acidity. However, although the system showed a significant acid-responsive increase in protein expression, some residual expression was observed under non-acidic conditions (pH 7.4). Further optimization will therefore be important for applications requiring a large dynamic range. Increasing the pH-responsive ssDNA-to-trigger ratio from 3:1 to 5:1 reduced background expression at pH 7.4, but also substantially decreased expression under acidic conditions, indicating a trade-off between suppressing leakage and maintaining activated output (**Supplementary Fig. 18**). Since the molecular origin of the residual expression was not fully resolved in this study, future optimization will require further investigation of potential contributions from incomplete trigger sequestration or spontaneous trigger dissociation. Further tuning of the pH-responsive strand–trigger interaction or incorporation of additional regulatory mechanisms may reduce basal expression and improve the dynamic range. For example, combinatorial gene-regulation strategies such as AND-gate logic^[21]^ could further minimize leakage while improving response specificity. Such approaches may enable synthetic cells to respond autonomously to more narrowly defined conditions—for example, by producing anti-inflammatory proteins only when both acidic pH and histamine^[10]^ are present.

A central challenge in this work was coupling the pH-responsive ssDNA to the toehold switch, since the pH required for trigger strand release was lower than the range where cell-free protein synthesis was active. Through iterative design, we found that the annealing length between the pH-responsive and trigger strands critically determines the trigger-release transition midpoint (pH1/2). Shortening the annealing length increased the pH1/2, enabling acid-responsive protein synthesis within the operational pH range of cell-free expression. Although previous studies have tuned the pH responsiveness of Hoogsteen interactions within the triplex-forming ssDNA itself,^[23,32]^ this work is the first to establish a quantitative design rule for controlling the transition midpoint governing hybridization and detachment between pH-responsive and complementary ssDNAs. This finding should facilitate more precise integration of pH-responsive DNA with other nucleic acid–based technologies.

Building on this design principle, our system could be further extended by reconfiguring the interactions. For example, the pH-responsive ssDNA can be designed to directly complement the toehold switch sequence so that, under acidic conditions, its detachment restores the riboregulator hairpin and suppresses protein translation. A three-way junction (3WJ) repressor^[46]^ that forms a hairpin around the RBS can instead be hybridized with pH-responsive ssDNA to achieve a similar outcome. Alternatively, embedding pH-responsive motifs directly into the toehold switch RNA could enable self-regulated, pH-dependent control of translation. Such systems could also be applicable in living cells—for instance, activating proton pump expression under acidic stress to improve cell viability.^[47,48]^

We also demonstrated that embedding pH-responsive synthetic cells into a hydrogel provides a strategy for developing pH-responsive, protein-producing biomaterials. Unlike conventional pH-responsive hydrogels that only release preloaded payloads, this system enables *in situ* protein synthesis in response to acidity. Overall, this work establishes a molecular strategy for programming synthetic cells to convert environmental acidity into nucleic acid information that drives cell-free protein synthesis. By linking acid sensing to gene expression, it expands the repertoire of signals that synthetic cells can interpret and enables direct responses to critical environmental cues, thereby broadening their potential applications in responsive biomaterials, biosensing, and controlled delivery.

## Supporting Information

Supporting Information is available from the Wiley Online Library or from the author.

## Acknowledgements

We thank Hossein Moghimianavval (Boston University, USA) for helpful discussions. We thank Seoyoung Kang (University of Michigan, USA) for help with experiments. S.-W.H. acknowledges the National Institutes of Health (NIH) Ruth L. Kirschstein Predoctoral Fellowship (1F31HL170510). A.P.L. acknowledges support by NIH R01 (EB030031 and R01GM163198) and R21 (AI173559) and National Science Foundation (2452482). A.A.G acknowledges support by NIH U01 (1U01AI148319), NIH R01 (1R01EB031893), NIH Transformative R01 (R01EB037112), and Boston University startup funds. The content is solely the responsibility of the authors and does not necessarily represent the official views of the National Institutes of Health. Any opinion, findings, and conclusions or recommendations expressed in this material are those of the author(s) and do not necessarily reflect the views of the National Science Foundation.

## Data Availability Statement

The data that support the findings of this study are available from the corresponding author upon reasonable request.

