## Supplementary Information for "pH-Responsive Synthetic Cells for Programmable *in Situ* Protein Synthesis and Release"

### Experimental Section

#### Materials

1-palmitoyl-2-oleoyl-glycero-3-phosphocholine (16:0-18:1 PC, POPC), 1,2-dioleoyl-sn-glycero-3-phosphoethanolamine-N-(lissamine rhodamine B sulfonyl (18:1 Liss Rhod PE), and 1-palmitoyl-2-oleoyl-sn-glycero-3-phospho-(1'-rac-glycerol) (16:0-18:1 PG, POPG) were purchased from Avanti Polar Lipids (Alabaster, AL). PUREfrex®2.0 (GeneFrontier) was purchased from Cosmo Bio USA. RNase Inhibitor, Murine (M0314S), Q5® High-Fidelity DNA Polymerase (M0491S), Deoxynucleotide (dNTP) Solution Mix (N0447S) were purchased from New England Biolabs. Sodium alginate (#I-1G, high stiffness gelation type, from Kimica [Tokyo, Japan]) was a gift from Prof. Sungmin Nam (University of Michigan). Nano-Glo® luciferase assay substrate (Cat# N1110) was purchased from Promega (Madison, WI, USA). Ni-charged magnetic beads (Cat# L00295) were purchased from GenScript (Piscataway, NJ, USA). Thermo Scientific™ Nunc 96-well optical-bottom plate (#165305) was purchased from Thermo Fisher Scientific Co. (Waltham, MA, USA). Corning® Elplasia® 384 well microplate (#4447), Corning® CellBIND® 384 well microplate (#CLS3770), Corning® 384 well low volume microplate (#CLS3540), cholesterol, light mineral oil (Cat# M5904), OptiPrep™ density gradient medium, Gramicidin A (#50845) from *Bacillus brevis*, 8-hydroxypyrene-1, 3, 6-trisulphonic acid trisodium salt (HPTS), Triton™ X-100 and all other chemicals were purchased from Millipore-Sigma (St. Louis, MO, USA) unless otherwise specified. AF647 and cholesterol-modified/unmodified DNA oligonucleotides, DNA gene fragments, and primers for cloning were synthesized by Eurofins. CPP-sfGFP-His and sfGFP-His linear DNA fragments were synthesized by Twist Bioscience. All DNA sequences can be found in Supplementary Notes.

#### Vesicle generation

Vesicles were generated using the inverted emulsion method[1]. First, lipid-in-oil dispersion was made by adding 41.0 µL of 25 mg/mL POPC, 1.16 µL of 50 mg/mL cholesterol, and 1.95 µL of 1 mg/mL rhodamine-PE (all stock solutions in chloroform) in a 20 mL glass vial to achieve 89.9/10/0.1 mol% of POPC/cholesterol/Rhod-PE. For cell-penetrating peptide-containing experiments, 29.6 µL of 25 mg/mL POPC, 28.9 µL of 10 mg/mL POPG, 1.16 µL of 50 mg/mL cholesterol, and 1.95 µL of 1 mg/mL rhodamine-PE were used instead, to achieve 64.9/25/10/0.1 mol% of POPC/POPG/cholesterol/Rhod-PE. The chloroform was removed by gentle argon flow, then further dried in a vacuum desiccator for 30 minutes. Lipids were rehydrated in 3 mL mineral oil to a final total lipid concentration in oil of 0.5 mM. The vial was then sealed, bath sonicated for 20 minutes, incubated at 60 °C for 1 hour, vortexed for 2 minutes, and bath sonicated again for 20 minutes to ensure complete dispersion. Next, the oil−water interface was formed by gently layering 300 µL of the lipid-in-oil dispersion over 400 µL of vesicle outer solution in a 1.5 mL tube. The outer solution (approximately 200 mM Tris and 680 mM HEPES, pH 7.4) was prepared by diluting a 0.5 M Tris/1.7 M HEPES stock solution (pH 7.4) with distilled water to match the osmolarity of the inner solution. The mixture was then incubated at room temperature for 30–60 minutes. During the incubation, the inner encapsulation solution was prepared in a separate 1.5 mL tube. For protein-expressing synthetic cells, a total of 10–12 µL inner solution contained PUREfrex®2.0, supplemented with 2 nM plasmid or linear DNA, 1,000 units/mL murine RNase inhibitor, 5 v/v% OptiPrep™ density solution, 1.2 µM trigger ssDNA or 1.2 µM trigger ssDNA hybridized with pH-responsive ssDNA. 600 µL of lipid-in-oil dispersion was added to the inner solution, and the mixture was thoroughly pipetted up and down for 3 min to produce water-in-oil monolayer emulsion droplets. The droplets were then added on top of the pre-formed oil–water interface and centrifuged at 2,500 g for 15 min at 15 °C. After centrifugation, the oil phase was carefully removed using the pipette, and the vesicles were collected into a 0.2 mL PCR tube using a fresh pipette tip, followed by gentle resuspension.

**pH-dependent hybridization/detachment test of pH-responsive ssDNA and trigger ssDNA**

AF647-labeled pH-responsive ssDNA and cholesterol-labeled trigger ssDNA were annealed at a 1:1 molar ratio using Duplex Buffer (Integrated DNA Technologies, IDT), from the respective stocks in 100 µM. Annealing was performed in a thermocycler using the following protocol: 5 minutes at 95 °C, followed by stepwise cooling—2 minutes per degree until the melting temperature (Tm) of the constructs, then 30 minutes per degree down to Tm − 5 °C, and finally 2 minutes per degree until reaching 25 °C. The annealed constructs were diluted to 1.2 µM and used as the inner solution for vesicle generation, supplemented with 5% v/v OptiPrep™ density solution. The outer solution was prepared by diluting a stock solution consisting of 0.5 M Tris and 1.7 M HEPES (pH 7.4). Vesicles were generated and subsequently added to wells of a 96-well plate containing various pH conditions. Gramicidin A was added to achieve a final concentration of 10 µg/mL to allow proton channel formation in the membrane. Vesicles were incubated overnight at room temperature, then imaged using fluorescence microscopy.

**Designing and screening of toehold switch sensors**

Toehold switches were designed to be activated by a single-stranded DNA (ssDNA) trigger that is partially complementary to a pH-responsive ssDNA. Switches matching these criteria were identified using a NUPACK-based selection algorithm[2,3]. The design of these candidate devices specifically targeted a 26-nucleotide continuous segment of the trigger ssDNA. For each putative toehold switch, multiple ensemble defect levels were computed to assess the deviation from its ideal secondary structure. Furthermore, the binding affinity between the target (RNA version) and toehold switch RNAs was quantified by calculating the equilibrium fraction of target/toehold switch complexes under equimolar conditions. This process generated 25 candidate toehold switches, which varied in toehold region and stem length. Initial screening of these candidates was conducted using a pHtdGFP output in a cell-free reaction based on their ON/OFF ratio. The best-performing sensors were subsequently identified with NLuc expression inside the synthetic cells under different pH conditions.

**Cloning and preparing DNA constructs**

DNA plasmids encoding pHtdGFP and NLuc were constructed via standard cloning methods. Synthetic gene fragments containing pHtdGFP and NLuc coding sequences were purchased, and the backbone fragments were PCR-amplified from a modified pET-28b vector (with the LacO operator removed), purified, and assembled using Gibson Assembly. For experiments involving CPPs and toehold switches with sfGFP, linear DNA fragments were directly synthesized rather than cloned into plasmids. Repeated attempts to assemble a plasmid construct containing both toehold switch and CPP elements were unsuccessful, likely due to toxicity that impeded propagation in *E. coli*.

**Bulk cell-free protein expression**

Cell-free protein synthesis was carried out using the PUREfrex® 2.0 kit (GeneFrontier) according to the manufacturer's instructions. Each reaction (5 µL total volume) contained either plasmid DNA or linear DNA template (final concentration: 2 nM), murine RNase inhibitor (1,000 units/mL), and either trigger ssDNA or an annealed construct consisting of trigger ssDNA hybridized to the pH-responsive ssDNA. For reactions with plasmid DNA (pHtdGFP and NLuc), the trigger ssDNA or annealed construct was added at a final concentration of 1.2 µM; for reactions with linear DNA templates (CPP-sfGFP), the final concentration was 4.8 µM. Reactions were transferred to a low-volume 384-well microplate, sealed, and incubated at 37 °C for 16 h in a Synergy H1 plate reader (BioTek). pHtdGFP fluorescence was measured every 10 min ($\lambda$ex = 488 nm, $\lambda$em = 528 nm).

**Confocal imaging**

Images were acquired using an inverted microscope (IX-81, Olympus) equipped with a spinning-disk confocal scanner (CSU-X1, Yokogawa), an oil-immersion Plan-Apochromat 60×/1.4 NA objective (Olympus), an iXON3 EMCCD camera (Andor Technology), and a National Instruments DAQ-MX–controlled laser system (Solamere Technology). Image acquisition was performed using MetaMorph software (Molecular Devices). Fluorescence images of pHtdGFP and sfGFP were obtained using 488-nm excitation, and Rhod-PE incorporated in the vesicle membrane was imaged using 561-nm excitation. All images were analyzed using ImageJ (NIH).

**Time-lapse fluorescence imaging of pHtdGFP expression**

Synthetic cells were imaged using an Agilent BioTek Cytation 5 Cell Imaging Multi-Mode Reader (20× objective, 1 h intervals, 20 h, 37 °C). Rhod-PE-labeled vesicle membranes were imaged in the RFP channel (531/593 nm), and pHtdGFP fluorescence in the GFP channel (469/525 nm). Using ImageJ, the mean fluorescence intensity of each vesicle was measured within a circular ROI (~2/3 of the vesicle diameter) and background-subtracted at each time point. Intensities were then normalized to each vesicle's maximal value, and the normalized time courses were fitted with a Boltzmann function to determine the response midpoint (t50) and the time to reach 90% of the fitted maximal response (t90).

**Protein expression in synthetic cells under different pH conditions**

Synthetic cells were prepared by encapsulating PUREfrex® 2.0 cell-free expression reactions containing either toehold switch–containing plasmid DNA or linear DNA and trigger ssDNA (either free or pre-annealed with the pH-responsive ssDNA) inside vesicles, as described above. The vesicles were collected and resuspended, and equal volumes (5 v/v%) of the vesicles were transferred into wells of a 384-well plate containing solutions at different pH values (pH 7.4, 7.0, 6.5, 6.0, and 5.5). Each external solution was supplemented with gramicidin A (added from a 10 mg/ml DMSO stock to a final concentration of 10 µg/mL) and the small-molecule components required for cell-free protein synthesis, including buffering salts, magnesium acetate, potassium glutamate, creatine phosphate, and nucleoside triphosphates[1,4]. The plate was then sealed and incubated at 37 °C for 20 h. Following incubation, to minimize pH-dependent differences in fluorescence (pHtdGFP/sfGFP) or NanoLuc activity and allow comparison based on the amount of protein produced, all samples were adjusted to neutral pH at equal final volumes. Confocal imaging was then performed, or a membrane-permeable NanoLuc substrate was added, and luminescence was measured using a Synergy H1 plate reader (BioTek Instruments).

#### Quantification of AF647 fluorescence ratio between the lumen and membrane of vesicles

AF647 fluorescence intensity profiles across individual vesicles were obtained using ImageJ (see **Fig. 1c**). Representative examples of the line-profile analysis are shown in **Supplementary Fig. 7**. The two membrane positions were identified from the membrane-associated fluorescence peaks or, when peaks were obscured by high luminal fluorescence, from the steepest intensity transitions at the membrane boundaries. Membrane intensity was the mean of these two positions. Lumen intensity was the mean of three points 5–7 pixels inward from each boundary (six points total), minimizing membrane contribution. Background intensity was the mean of three points 10–12 pixels outward from each boundary (six points total) and was subtracted from both lumen and membrane values before calculating the ($F_{Lumen}/F_{Membrane}$) ratio. A uniform scaling factor of 0.8 was applied so that the fully lumen-localized reference condition yielded a ratio of approximately 1. Only isolated vesicles with clearly defined membrane boundaries and without visible membrane aggregates were included in this quantitative analysis.

#### pH-responsive synthetic cell embedment in hydrogel

Alginate solutions (1.5 w/v%) were prepared at two different pH values (resulting pH to be pH 7.4 and 6.5 at 37 °C). pH-responsive synthetic cells were prepared as described above and pre-incubated with gramicidin A (20 µg/mL) for 15 min on ice. The alginate solution and synthetic cells were then gently mixed to obtain a final alginate concentration of 0.8%. Next, 10 µL of the mixture was transferred into a 1.5 mL microcentrifuge tube using a cut pipette tip to avoid shear. To initiate gelation, 80 µL of 50 mM $CaCl_2$ (prepared in the corresponding pH-matched buffer) was carefully added on top of the alginate mixture without disturbing the surface using a cut pipette tip, and the sample was allowed to crosslink for 5 min at room temperature. The resulting hydrogel was washed twice with the respective $Ca^{2+}$-free pH buffer and transferred to a 96-well plate. Each well was supplemented with 40 µL of the corresponding pH buffer containing the small-molecule components required for cell-free protein synthesis. The plate was sealed and incubated at 37 °C for 20 h.

#### Capturing and quantifying released sfGFP using Ni-NTA–coated magnetic beads

Synthetic cells expressing CPP-tagged or untagged sfGFP-His were prepared and incubated at 37 °C for 20 h as described above. Ni-NTA–coated magnetic beads (GenScript) were washed three times with pH 7.4 buffer (200 mM Tris–680 mM HEPES, adjusted to match the osmolarity of the external solution). An equal volume of washed beads was then added to each well, followed by incubation for 1 h at room temperature prior to imaging. For control experiments performed at pH 7.4, comparing sfGFP without and with CPP (−CPP and +CPP), washed Ni-NTA beads were added directly after the incubation. In contrast, for experiments comparing pH-responsive synthetic cells at pH 7.4 and pH 6.0 using the annealed construct, Ni–His binding could not occur at pH 6.0. Therefore, after 20 h incubation, the vesicle-containing solutions were adjusted to pH 7.4 by diluting to equal final volumes before adding the washed Ni-NTA beads. The samples were then incubated for 1 h at room temperature, and sfGFP fluorescence on the bead surfaces was imaged.

For fluorescence quantification, the dark central region of each bead was masked using the ImageJ Threshold tool (see **Supplementary Fig. 16**). The mean pixel intensity of the remaining fluorescent bead boundary region was measured, and the mean background pixel intensity was subtracted to obtain the corrected sfGFP signal.

**pH measurement and limitations**

pH values were measured using a Thermo Scientific™ Orion™ Star A111 pH meter equipped with Thermo Scientific™ Orion™ PerpHecT™ ROSS™ pH microelectrode. For experiments requiring incubation at a defined pH at 37 °C, samples were equilibrated at 37 °C in a water bath before pH measurement to account for temperature-dependent changes in pH. The meter was calibrated before each use using standard pH 4, 7, and 10 buffers. The microelectrode was sufficiently immersed in the sample, and pH was recorded after the reading had stabilized. To validate the small-volume measurements, five representative solutions were independently measured using a conventional glass pH electrode (edge® Dedicated pH/ORP Meter, Hanna Instruments) and compared with measurements obtained using the microelectrode setup. The conventional electrode yielded pH values of 6.09, 7.42, 7.80, 7.82, and 9.02, compared with 6.07, 7.44, 7.80, 7.86, and 9.06, respectively, using the microelectrode, corresponding to a maximum difference of 0.04 pH units. For the hydrogel experiments, pH could not be measured directly after $Ca^{2+}$-mediated crosslinking; therefore, the reported values represent the pH measured before crosslinking. To minimize potential pH changes associated with residual crosslinking solution, the gels were washed twice with $Ca^{2+}$-free solution and incubated in solutions adjusted to the intended pH.

**Statistical analysis**

Data processing, including normalization and exclusion criteria where applicable, is described in the corresponding experimental methods and figure captions. Data presentation and sample sizes ($n$), including whether $n$ represents individual vesicles/beads or independent experimental replicates, are specified in each legend. For two-group comparisons, an F-test was first performed to assess equality of variances, followed by a two-tailed Student's $t$-test or Welch's $t$-test, as appropriate. One-way ANOVA was followed by Tukey's *post hoc* test, whereas two-way ANOVA was followed by Sidak–Holm multiple-comparisons tests. The $t$-tests and F-tests were performed using Microsoft Excel, and all ANOVA analyses were performed using OriginPro. A significance level of $\alpha = 0.05$ was used. Statistical significance was indicated as follows: **** $p < 0.0001$, *** $p < 0.001$, ** $p < 0.01$, * $p < 0.05$.

### Supplementary Figures

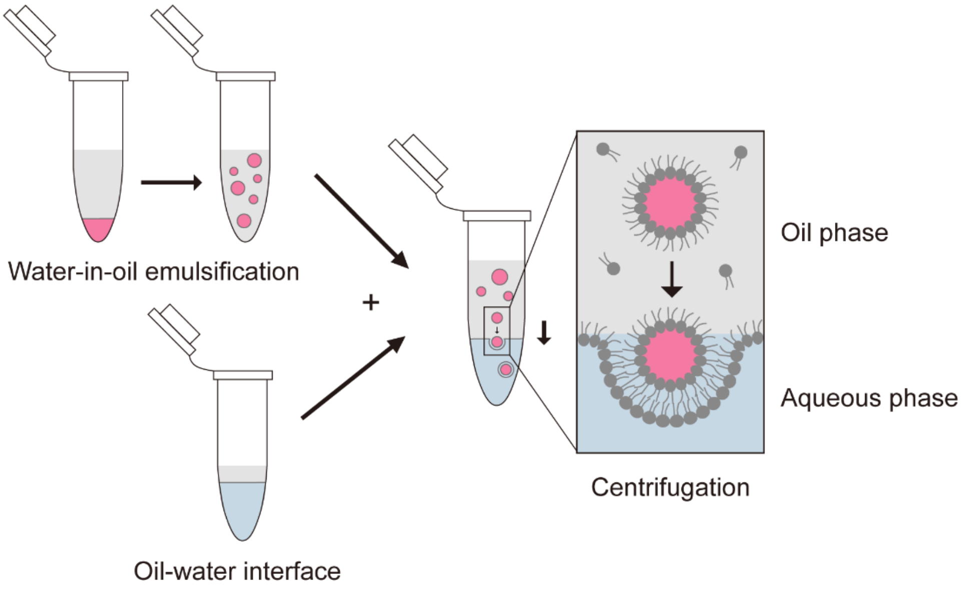


**Supplementary Fig. 1. Schematic illustration of vesicle generation using the inverted emulsion method.** Lipid-coated water-in-oil droplets are driven through a lipid monolayer formed at the oil–water interface by centrifugal force. During passage through the interface, the droplets acquire a second lipid monolayer through lipid–lipid interactions, including van der Waals attraction, resulting in the formation of lipid bilayer vesicles in the aqueous phase.

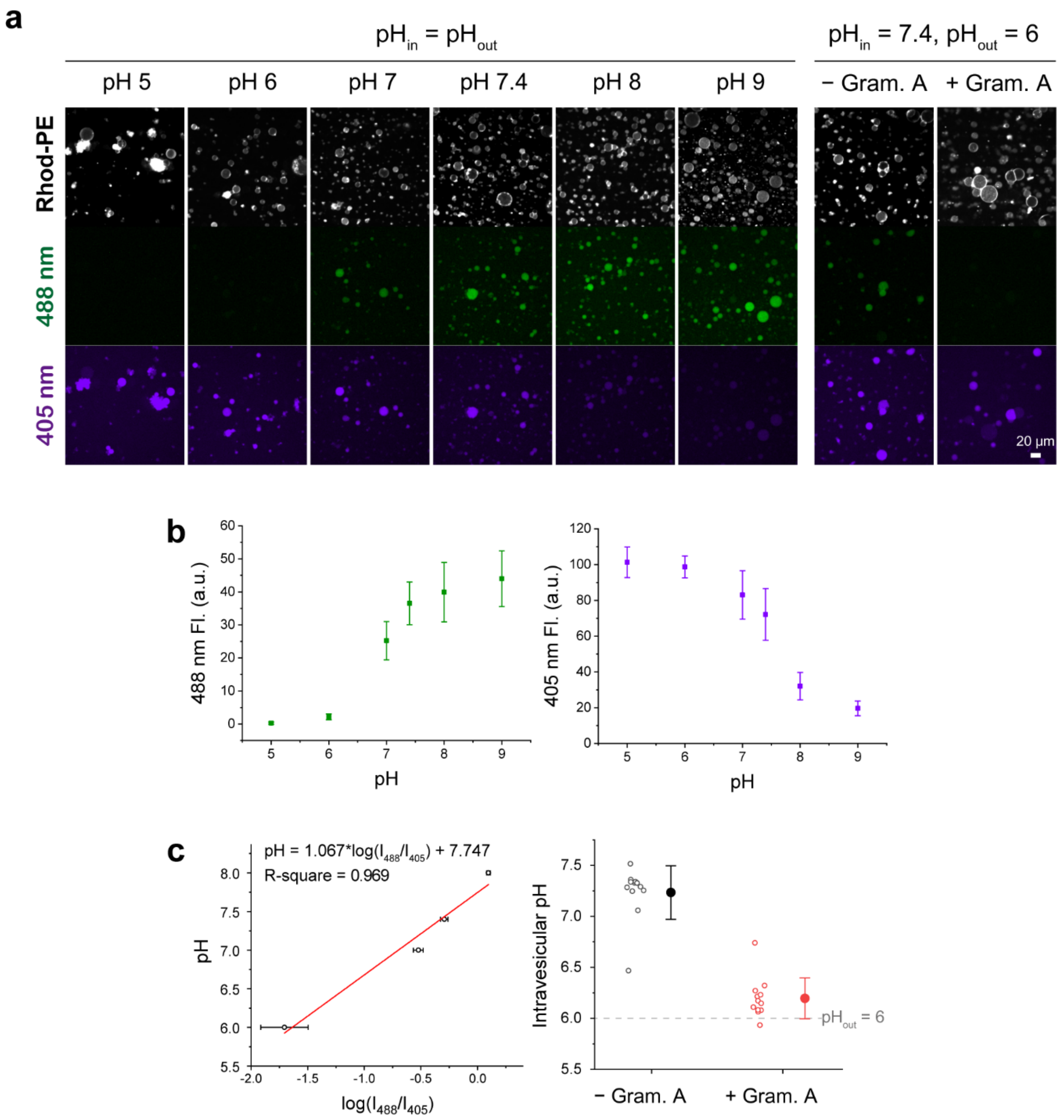


**Supplementary Fig. 2. Direct measurement of intravesicular pH demonstrates gramicidin A–mediated acidification of synthetic cells. a,** Representative fluorescence images of synthetic cells encapsulating the ratiometric pH indicator HPTS. For calibration, vesicles were prepared with matched internal and external pH (*i.e.*, $pH_{in} = pH_{out}$) ranging from pH 5 to 9. Rhod-PE indicates the vesicle membrane, and HPTS was imaged with 488- and 405-nm excitation. To evaluate gramicidin A–mediated proton transport, vesicles initially at $pH_{in} = 7.4$ were exposed to $pH_{out} = 6.0$ for 14 h with or without gramicidin A. **b,** Mean HPTS intensities under 488-nm (left) and 405-nm (right) excitation across the pH-matched calibration conditions. Error bars are mean ± s.d. ($n = 12$ vesicles per condition). **c,** Left, calibration curve of pH versus $\log(I_{488}/I_{405})$, fitted by linear regression using the four conditions within the experimentally relevant range (pH 6.0, 7.0, 7.4, 8.0). Right, intravesicular pH calculated from this calibration curve for vesicles exposed to $pH_{out} = 6.0$. Without gramicidin A, the intravesicular pH remained near the initial value (7.2 ± 0.3); with gramicidin A, it dropped to 6.2 ± 0.2, close to the external pH. Each point represents an individual vesicle.

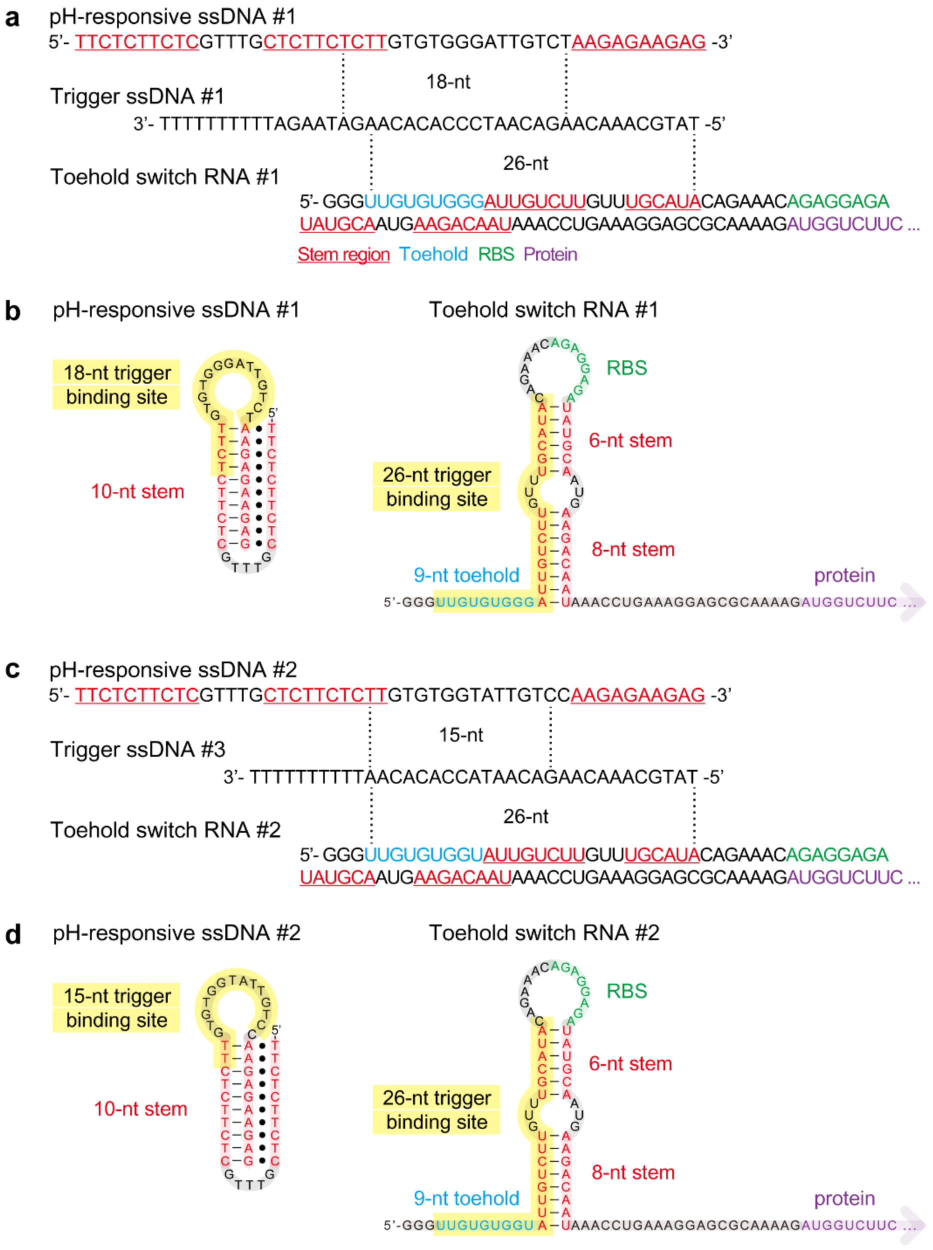


**Supplementary Fig. 3. Predicted binding regions and secondary structures of the pH-responsive ssDNA, trigger ssDNA, and toehold switch RNA. a**, Complementary regions (18 nt and 26 nt) between pH-responsive ssDNA #1, trigger ssDNA #1, and toehold switch RNA #1. Hairpin-forming stem domains are underlined. The trigger ssDNA is shown in the 3′→5′ orientation. See *Supplementary Table S1* for all other sequence variants (pH-responsive ssDNA #1–3, trigger ssDNA #1–3, and toehold switch #1–2). **b**, Predicted structures of the pH-responsive ssDNA #1 and toehold switch RNA #1 corresponding to panel **a**. **c**, Complementary regions (15 nt and 26 nt) between pH-responsive ssDNA #2, trigger ssDNA #3, and toehold switch RNA #2. **d**, Predicted secondary structures of the pH-responsive ssDNA #2 and toehold switch RNA #2 corresponding to panel **c**.

**a**

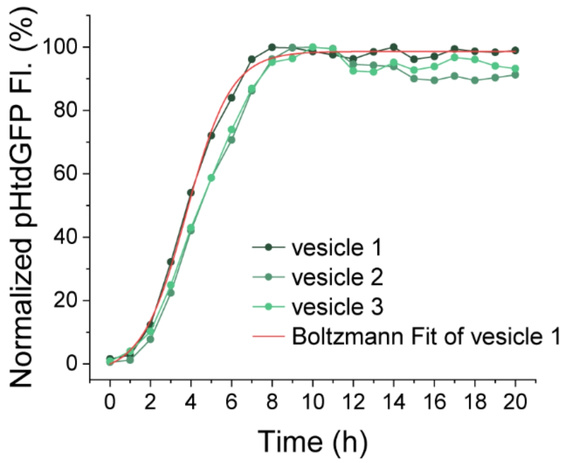


| Model | Boltzmann | | |
|---|---|---|---|
| Equation | y = A2 + (A1-A2)/(1 + exp((x-x0)/dx)) | | |
| Plot | vesicle 1 | vesicle 2 | vesicle 3 |
| A1 | -3.18968 ± 2.11165 | -2.50614 ± 4.06256 | -1.70742 ± 2.82207 |
| A2 | 98.6395 ± 0.54158 | 93.98801 ± 1.22721 | 95.61254 ± 0.80971 |
| x0 (= t50, h) | 3.77213 ± 0.08098 | 4.29807 ± 0.18034 | 4.28494 ± 0.126 |
| dx (h) | 1.09639 ± 0.06418 | 1.1414 ± 0.1472 | 1.20864 ± 0.10139 |
| t90 (h) | 6.18 | 6.81 | 6.94 |
| R-Square (COD) | 0.99747 | 0.98773 | 0.99471 |

**b**

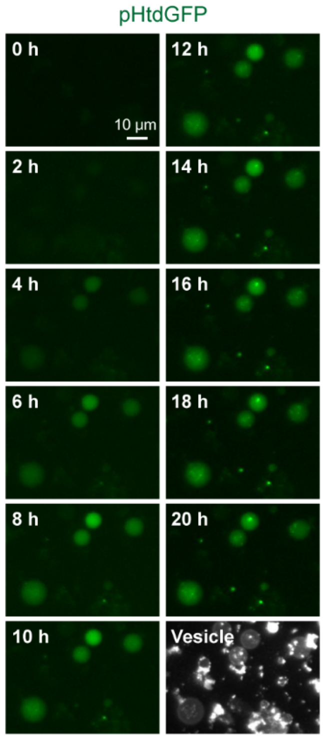


**Supplementary Fig. 4. Time-resolved pHtdGFP expression in acid-responsive synthetic cells. a**, Normalized pHtdGFP fluorescence of three individually tracked synthetic cells during incubation at pH 6.3. Fluorescence for each vesicle was normalized to the maximal (100%) fluorescence intensity. The red curve shows a representative Boltzmann fit for vesicle 1; fitting parameters for all three vesicles are summarized below the graph. The fitted response midpoint (t50) was 4.12 ± 0.30 h, and 90% of the fitted maximal response (t90) was reached at 6.64 ± 0.41 h (mean ± s.d., *n* = 3 individual vesicles). **b,** Representative time-lapse fluorescence images showing progressive pHtdGFP accumulation in synthetic cells from 0 to 20 h.

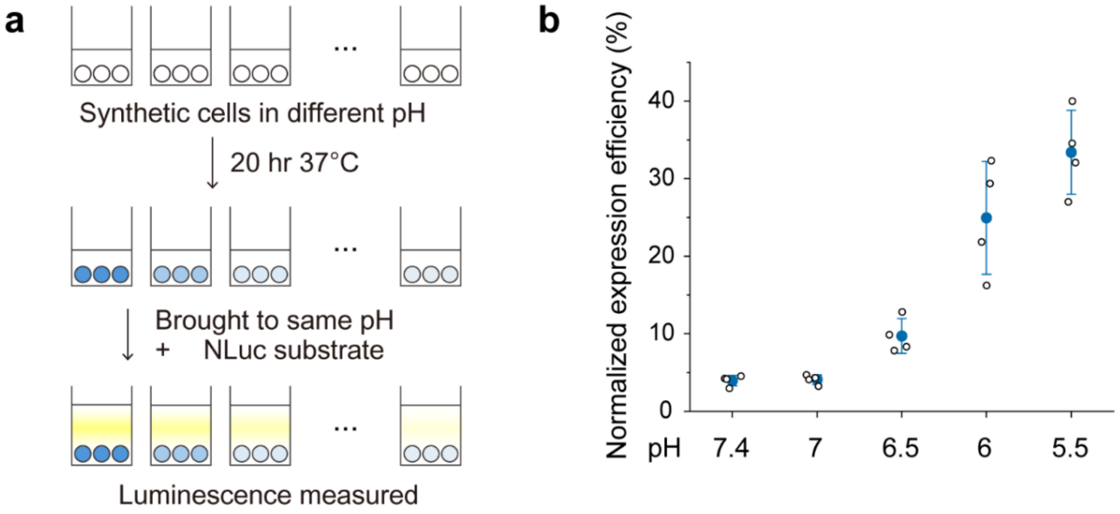


**Supplementary Fig. 5. Experimental procedure and normalization of NLuc expression in synthetic cells under different pH conditions. a,** Schematic of the experimental procedure. Synthetic cells encapsulating a toehold switch–containing NLuc plasmid, trigger ssDNA (either free or hybridized with pH-responsive ssDNA), and cell-free expression components were prepared and incubated with gramicidin A at 37 °C for 20 h under various pH conditions (pH 7.4, 7.0, 6.5, 6.0, and 5.5) in a 384-well plate. After incubation, all samples were adjusted to the same final pH of 7.3, followed by the addition of a membrane-permeable NLuc substrate furimazine, and luminescence was measured using a plate reader. **b,** NLuc expression from the pH-responsive system (**Fig. 2a**, right) normalized to the corresponding trigger control (**Fig. 2a**, left) at each pH. The trigger control condition, in which free trigger ssDNA was supplied, was defined as 100% at each pH to account for the intrinsic pH dependence of cell-free protein expression. Open circles represent individual experimental replicates, and filled circles with error bars indicate mean ± s.d. (*n* = 4 independent experiments).

**a**

| Model | Boltzmann | | | |
|---|---|---|---|---|
| Equation | y = A2 + (A1-A2)/(1 + exp((x-x0)/dx)) | | | |
| Plot | 18nt | 16nt | 15nt | 14nt |
| A1 | 0.95729 ± 0.1933 | 0.97718 ± 0.0353 | 0.98203 ± 0.0191 | 0.97206 ± 0.0182 |
| A2 | 0.12768 ± 0.0186 | 0.14756 ± 0.0181 | 0.11237 ± 0.0285 | 0.44771 ± 0.3149 |
| x0 | 6.16054 ± 0.4278 | 6.91184 ± 0.0371 | 7.21032 ± 0.0344 | 7.50959 ± 0.3135 |
| dx | 0.12 ± 0.14714 | 0.12 ± 0.04134 | 0.12 ± 0.02007 | 0.12 ± 0.23028 |
| Reduced Chi-Sq | 4.1523 | 1.64865 | 0.64217 | 0.73737 |
| R-Square (COD) | 0.90184 | 0.99646 | 0.99779 | 0.98333 |
| Adj. R-Square | 0.7546 | 0.99116 | 0.99447 | 0.95833 |

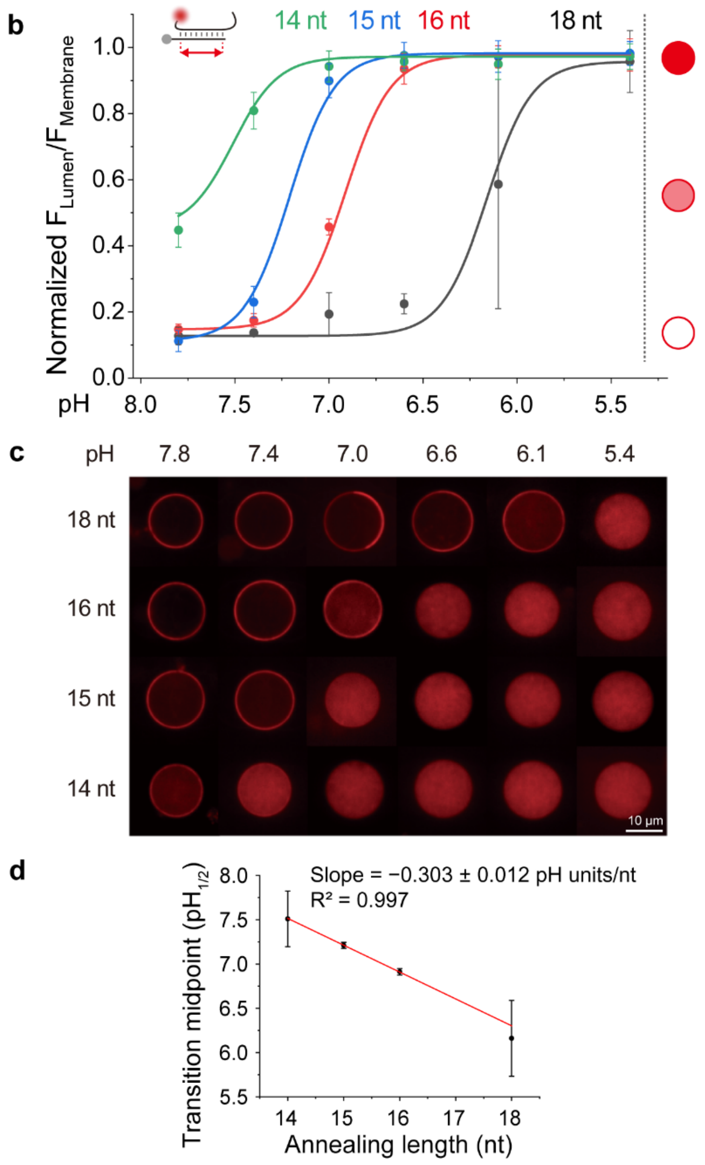


**Supplementary Fig. 6. Shorter annealing length increases the pH transition midpoint, indicating the transition from hybridized to detached pH-responsive ssDNA-trigger ssDNA. a,** Boltzmann fitting parameters for 18-, 16-, 15-, and 14-nt constructs, including the fitted transition midpoint ($pH_{1/2}$, $x_0$). **b**, Normalized fluorescence intensity ratio ($F_{Lumen}/F_{Membrane}$) as a function of pH. **c**, Representative AF647 fluorescence images for 14–18 nt constructs at pH 7.8–5.4. Panels **b** and **c** are also shown in **Fig. 2b**. **d,** Transition midpoint ($pH_{1/2}$) as a function of annealing length. Error bars represent the standard errors of $x_0$ from the Boltzmann fits. Weighted linear regression using inverse-variance weighting ($1/SE^2$) yielded a slope of −0.303 ± 0.012 pH units per nucleotide ($R^2$ = 0.997), indicating that shortening the annealing length by one nucleotide increased the transition midpoint by approximately 0.30 pH unit.

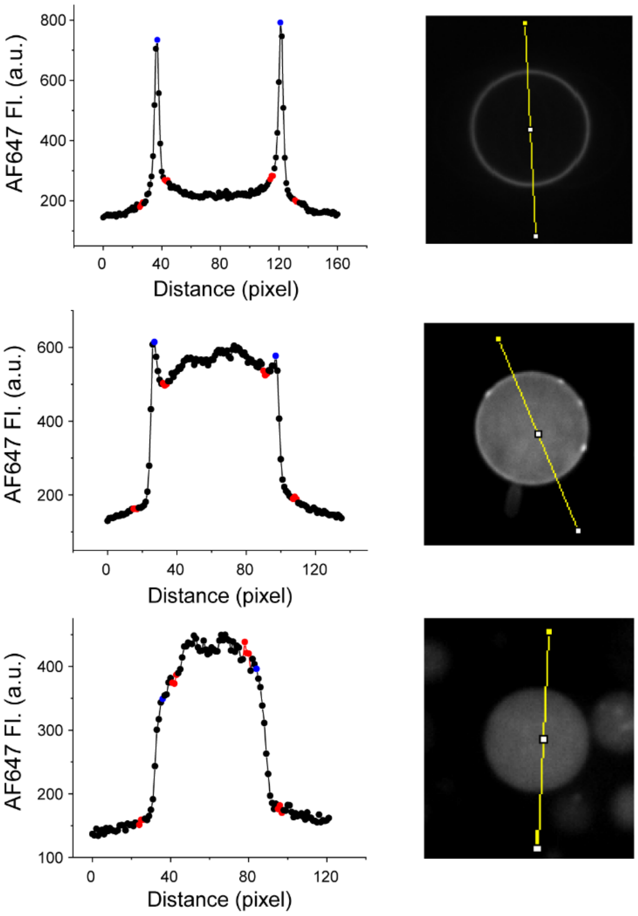


**Supplementary Fig. 7. Representative fluorescence intensity profiles used for AF647 localization analysis.** Representative AF647 fluorescence intensity profiles (left) and corresponding confocal images with the analyzed line (right), showing predominantly membrane-localized (top), intermediate (middle), and lumen-localized (bottom) fluorescence distributions. Blue points indicate the two membrane positions used to calculate membrane fluorescence. Red points indicate the positions used to calculate lumen and background fluorescence, located 5–7 pixels inward and 10–12 pixels outward from each membrane boundary, respectively. Background-subtracted $F_{Lumen}/F_{Membrane}$ ratios after application of the uniform 0.8 scaling factor were 0.113, 0.645, and 0.951 for the top, middle, and bottom examples, respectively.

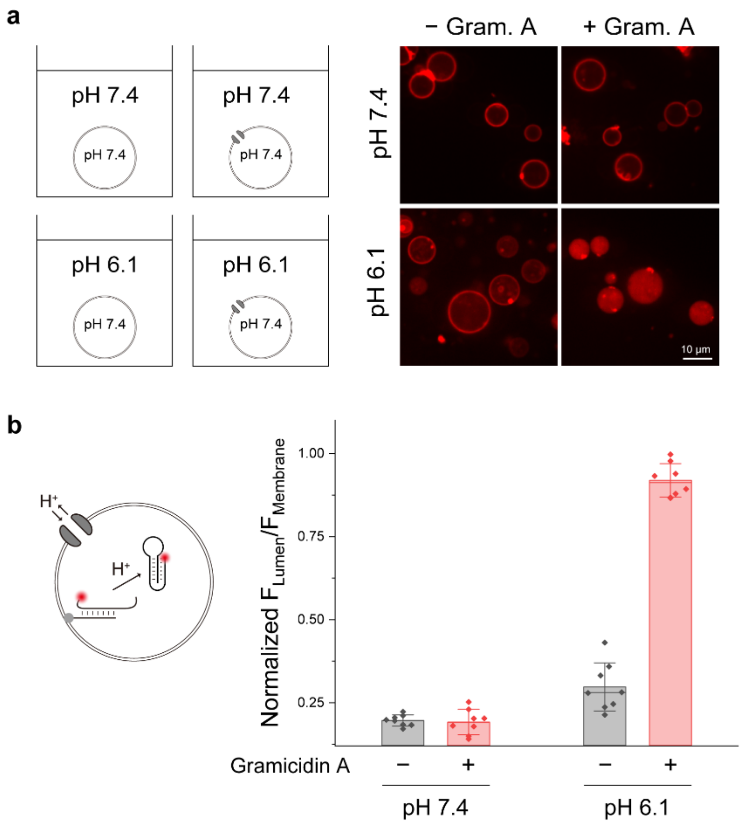


**Supplementary Fig. 8. Proton channel gramicidin A enables equilibration of external pH with the lumen of synthetic cells. a,** (Left) Schematic illustration of the experimental design used to evaluate whether gramicidin A is required for proton transport. Synthetic cells encapsulating Alexa Fluor 647 (AF647)–labeled pH-responsive ssDNA hybridized with cholesterol-tagged trigger ssDNA (16-nt annealing length; verified to exhibit complete luminal AF647 signal at pH 6.1, see **Fig. 2b**) were prepared at an internal pH of 7.4. The vesicles were then exposed to either pH 7.4 or 6.1 conditions for 18 h in the presence or absence of gramicidin A. The pH 6.1 solution contained 359 mM HEPES and 8.2 mM Tris, whereas the pH 7.4 solution contained 278 mM HEPES and 81.8 mM Tris. (Right) Representative fluorescence images of AF647 pH-responsive ssDNA. Vesicles containing gramicidin A and exposed to pH 6.1 displayed complete lumen fluorescence, indicating internal acidification. In contrast, vesicles without gramicidin A retained fluorescence predominantly at the membrane, suggesting insufficient acidification of the interior. **b,** Quantification of normalized fluorescence intensity ratio $F_{Lumen}/F_{Membrane}$. Error bars are mean ± s.d. ($n$ = 8 individual vesicles).

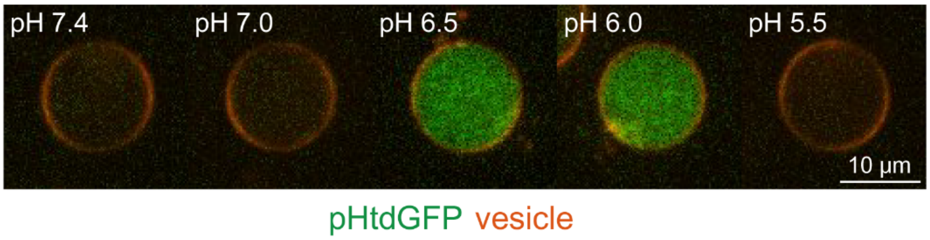


**Supplementary Fig. 9. Acid-responsive pHtdGFP expression inside synthetic cells for 20 hours with 15-nt annealing construct tested under pH 7.4–5.5.**

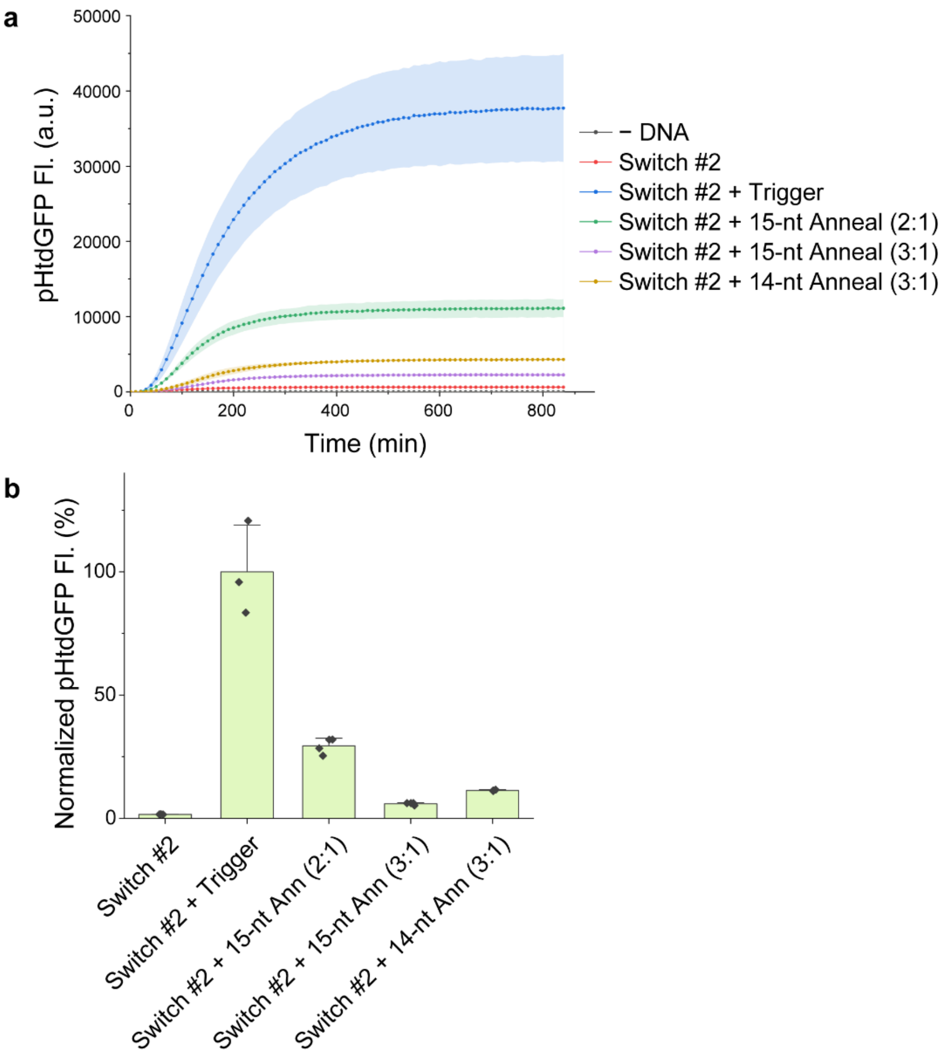


**Supplementary Fig. 10. pHtdGFP expression tested in bulk with reduced annealing lengths (15 nt and 14 nt). a,** Decreasing the annealing length from 18 nt to 15 nt caused an increase in leaky expression. This was reduced by increasing the pH-responsive:trigger ssDNA ratio from 2:1 to 3:1. **b**, Fluorescence is quantified at 14 h and normalized to the Switch #2 + Trigger condition. Error bars are mean ± s.d. (*n* = 3 independent samples).

**a**

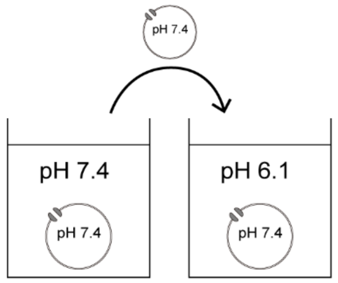


| Model | Exponential |
|---|---|
| Equation | y = y0 + A*exp(R0*x) |
| Plot | B |
| y0 | 0.95999 ± 0.01359 |
| A | -0.73564 ± 0.01393 |
| R0 | -0.12847 ± 0.01073 |
| Reduced Chi-Sqr | 0.00512 |
| R-Square (COD) | 0.99981 |
| Adj. R-Square | 0.99942 |

**b**

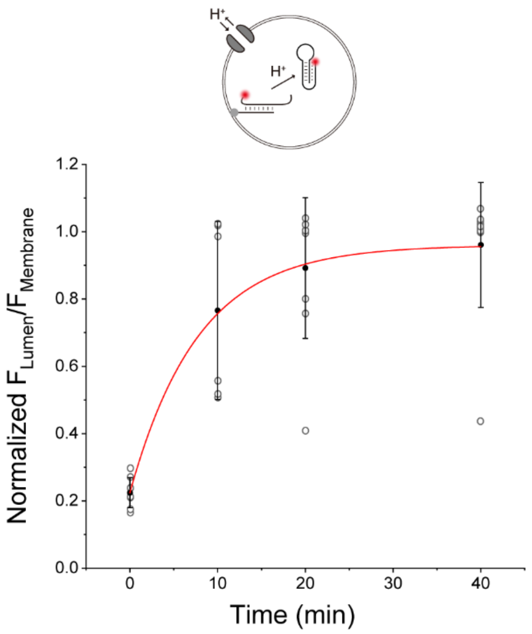


**Supplementary Fig. 11. Evaluation of internal acidification and trigger ssDNA release in pH-responsive synthetic cells. a,** Schematic illustration of experimental design. Synthetic cells encapsulating AF647–labeled pH-responsive ssDNA hybridized with cholesterol-tagged trigger ssDNA (16-nt annealing length; verified to exhibit complete luminal AF647 fluorescence at pH 6.1, see **Fig. 2b**) were prepared at an internal and external pH of 7.4 with gramicidin A and subsequently transferred to a pH 6.1 solution. The degree of ssDNA detachment or hybridization was monitored over time by measuring the fluorescence ratio $F_{Lumen}/F_{Membrane}$. **b**, Time-dependent changes in normalized fluorescence intensity ratio $F_{Lumen}/F_{Membrane}$. Approximately 73.5 % of strand detachment was observed within 10 min, and 90.7 % detachment was observed at 20 min. The red curve represents an exponential fit (see table in **a**). Error bars are mean ± s.d. ($n$ = 8 individual vesicles).

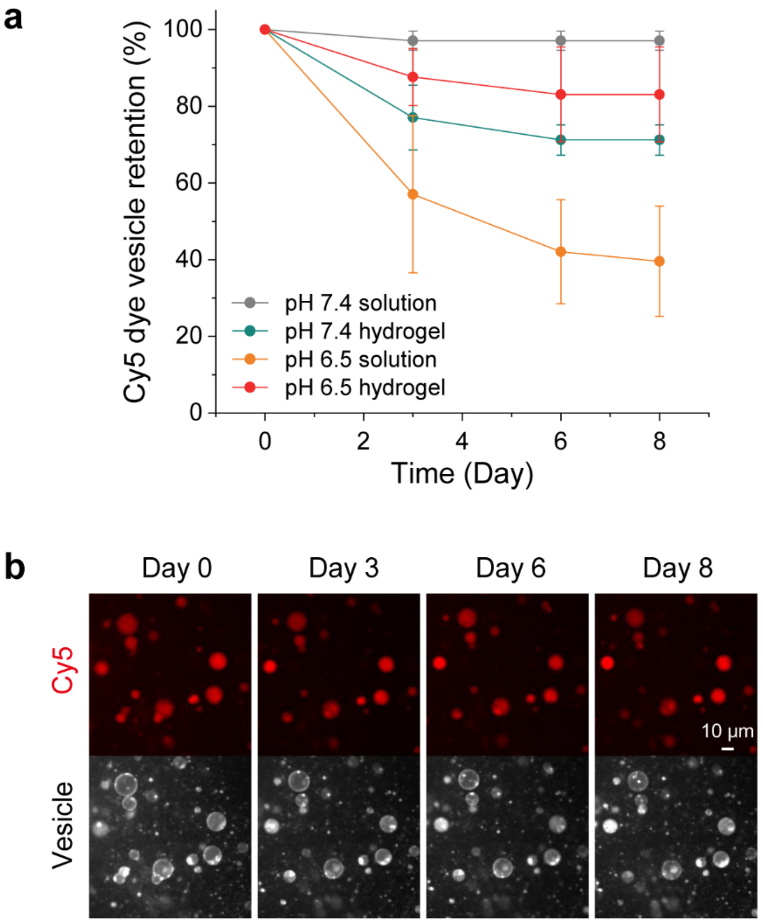


**Supplementary Fig. 12. Stability of synthetic cell vesicles in solution and alginate hydrogels over 8 days. a,** Vesicle retention in solution or alginate hydrogel at pH 7.4 and pH 6.5 over 8 days. The same vesicle populations were tracked at Days 0, 3, 6, and 8, and vesicles retaining detectable internal Cy5 fluorescence were counted. Vesicle numbers at each time point were normalized to the corresponding Day 0 value. Data are shown as mean ± s.d. ($n$ = 3–4 independently tracked imaging locations with total vesicle number tracked 58–63). **b,** Representative images of the same location in a pH 6.5 alginate hydrogel tracked over 8 days, showing internal Cy5 fluorescence (top) and corresponding vesicles (bottom).

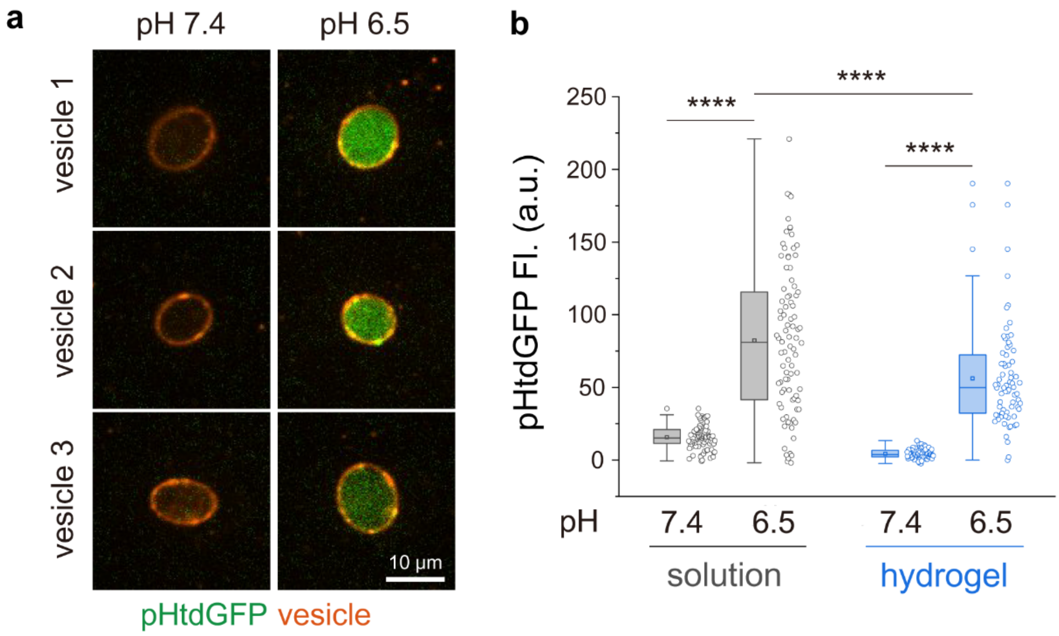


**Supplementary Fig. 13. Acid-responsive pHtdGFP expression in synthetic cells within pH-controlled hydrogels and comparison with synthetic cells in solution. a,** Representative confocal fluorescence images of synthetic cells embedded in hydrogels at pH 7.4 and pH 6.5. The images in the top row are the same as in **Fig. 3c**. Some vesicles exhibit non-spherical morphologies, likely due to mechanical deformation during handling, washing, and transfer of the crosslinked alginate hydrogels. **b,** Quantification of pHtdGFP fluorescence intensity in synthetic cells incubated in solution or embedded in hydrogels at pH 7.4 and pH 6.5. Individual vesicles are shown as data points, and box plots indicate the median and interquartile range. Statistical significance was assessed by two-way ANOVA with pH and embedding condition as factors, followed by Sidak–Holm multiple-comparisons test. $p$ values were $5.4 \times 10^{-36}$ (solution, pH 7.4 vs 6.5), $1.6 \times 10^{-20}$ (hydrogel, pH 7.4 vs 6.5), and $3.8 \times 10^{-7}$ (solution vs hydrogel at pH 6.5). ****$p < 0.0001$ ($n$ = 71–93 individual vesicles).

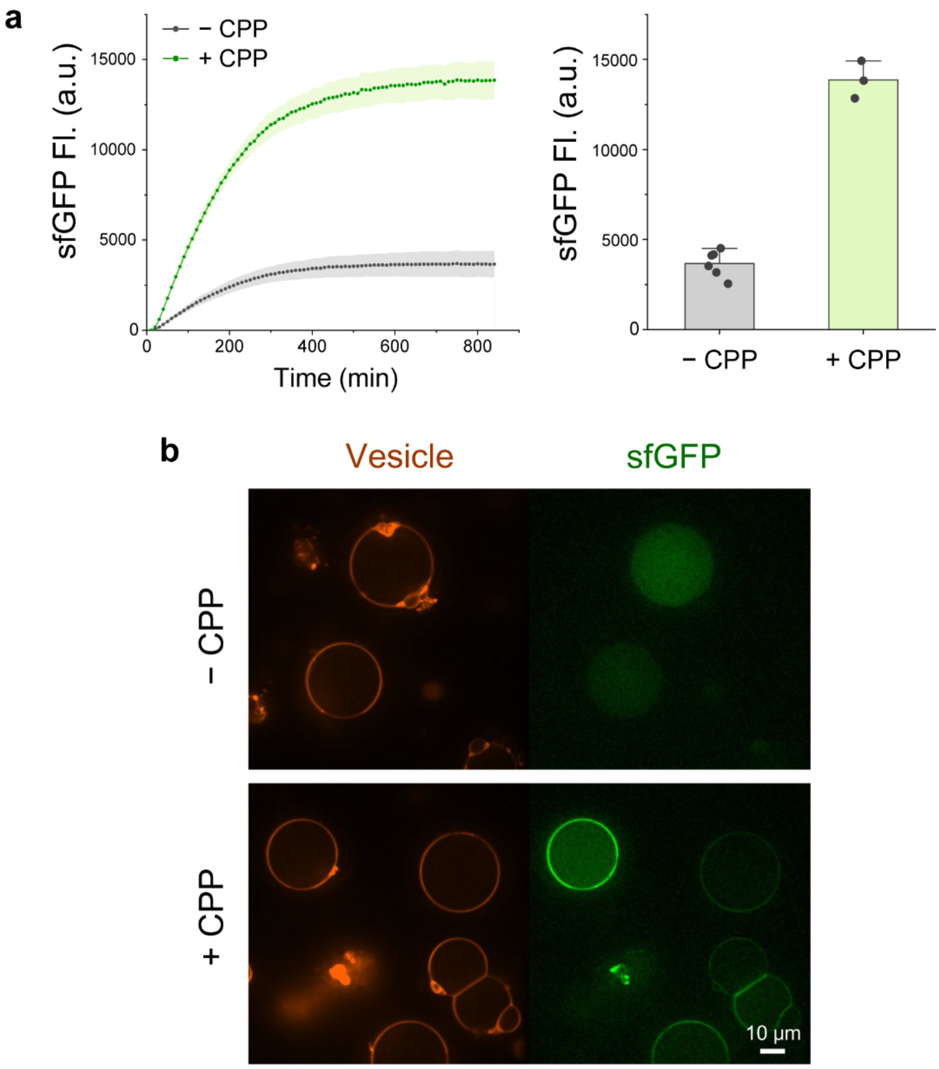


**Supplementary Fig. 14. Bulk expression and membrane localization of sfGFP-His and CPP-sfGFP-His. a,** Bulk cell-free expression of sfGFP-His (−CPP) and penetratin-fused sfGFP-His (+CPP), both using a 2 nM DNA template concentration. (Left) time-course measurements of sfGFP fluorescence during incubation at 37 °C for 14 h. (Right) endpoint sfGFP fluorescence measured at 14 h. Error bars are mean ± s.d. ($n$ = 3–6 independent cell-free reactions). **b,** Representative confocal fluorescence images of synthetic cells expressing sfGFP-His (top) and CPP-sfGFP-His (bottom). Synthetic cells expressing CPP-sfGFP-His exhibited fluorescence predominantly localized at the membrane, whereas those expressing sfGFP-His alone showed luminal fluorescence.

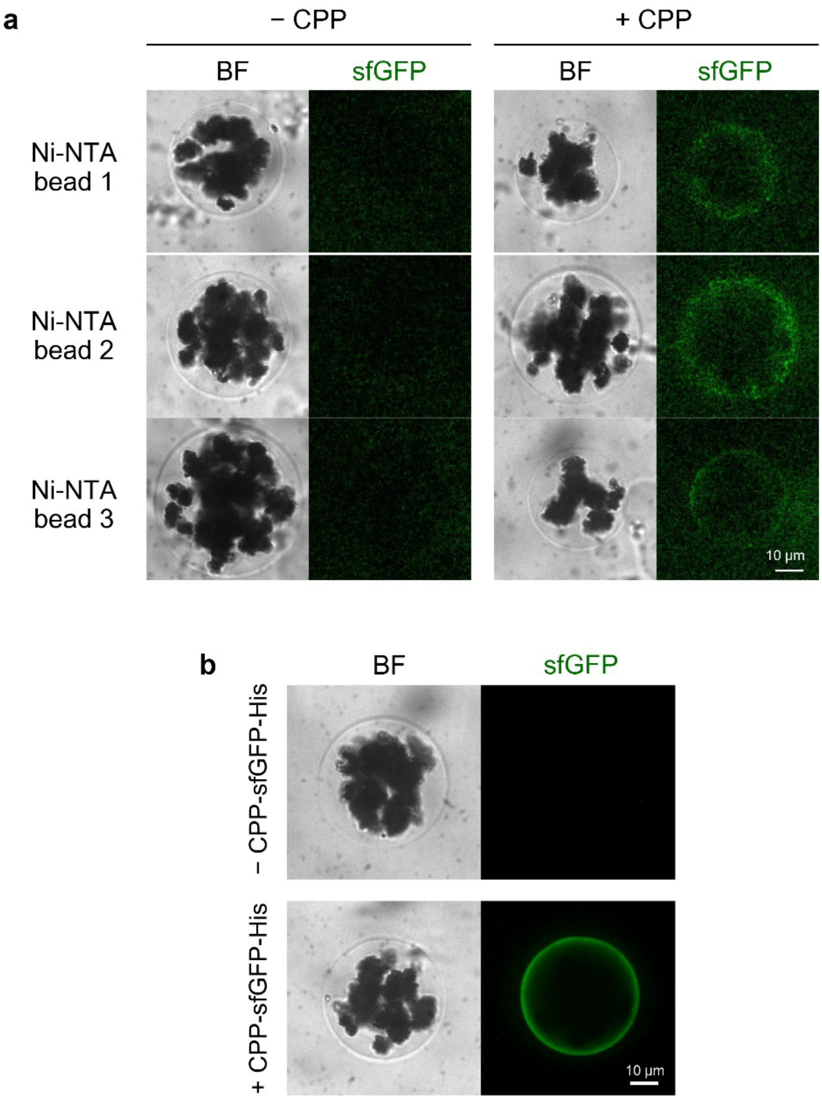


**Supplementary Fig. 15. Capture of sfGFP at Ni–NTA–coated magnetic bead surfaces. a**, Confocal fluorescence images of Ni-NTA-coated magnetic beads incubated with synthetic cells expressing sfGFP-His (without CPP) or CPP-sfGFP-His. sfGFP fluorescence was detected on bead surfaces only when incubated with CPP-sfGFP-His-expressing synthetic cells. The images in the top row are the same as in **Fig. 4b**. **b**, Control experiment showing Ni-NTA beads imaged without (top) or with (bottom) addition of bulk cell-free reaction expressing CPP-sfGFP-His.

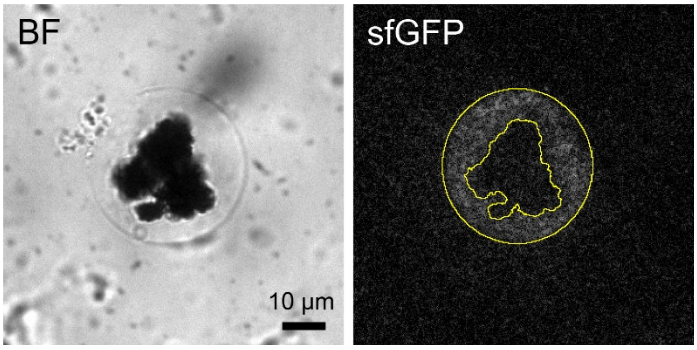


**Supplementary Fig. 16. Quantification of sfGFP fluorescence on Ni-NTA magnetic beads.** The dark core of each bead was masked using the Threshold tool in ImageJ (inner yellow contour). Fluorescence intensity was measured from the bead boundary region (the area between the inner and outer contours) and corrected by subtracting the background mean pixel intensity.

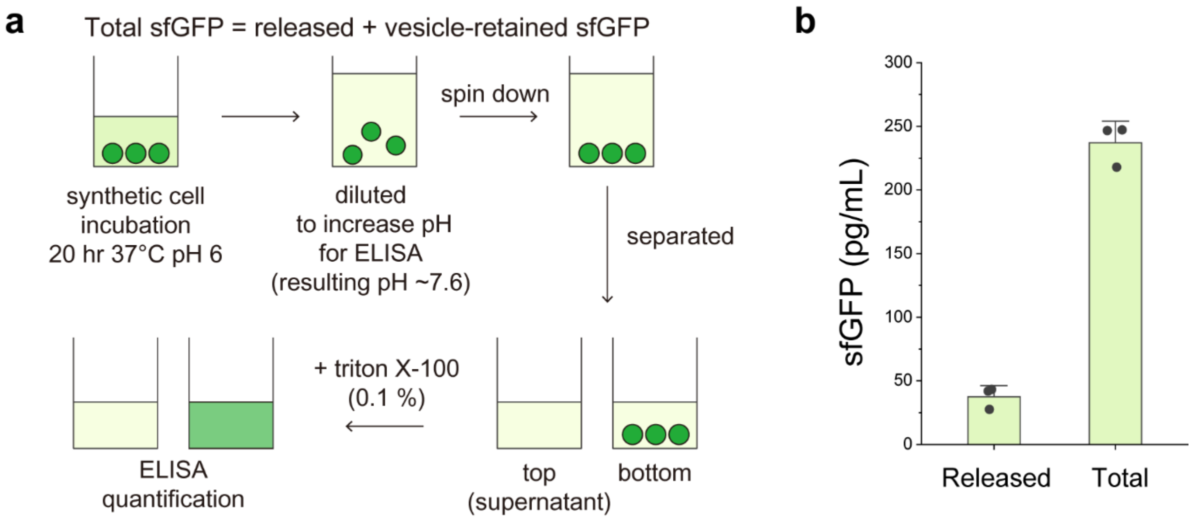


**Supplementary Fig. 17. Quantification of CPP-sfGFP-His release efficiency from synthetic cells. a,** Schematic of the procedure used to quantify released and total sfGFP. Synthetic cells were incubated for 20 h at 37 °C under acidic conditions (pH 6.0), followed by dilution to increase the pH to approximately 7.6 for ELISA analysis. After centrifugation, the upper half of the supernatant was collected and analyzed by ELISA. The remaining lower fraction, containing the synthetic cells, was lysed with 0.1% Triton X-100 and analyzed separately by ELISA, with the same Triton X-100 concentration also applied to the collected supernatant and sfGFP standards. The total amount of released sfGFP was estimated as twice the amount measured in the collected upper half, assuming that the intravesicular volume was negligible (4 µL vesicle suspension in a final volume of 100 µL). Total sfGFP was determined by summing the amounts measured in the collected supernatant and the Triton X-100-treated lower fraction. Release efficiency was calculated as the estimated released sfGFP amount divided by the total sfGFP amount. **b,** Released and total sfGFP concentrations determined by ELISA after 20 h. The calculated release efficiency was 15.7 ± 2.7%. Data are mean ± s.d. ($n$ = 3 independent samples).

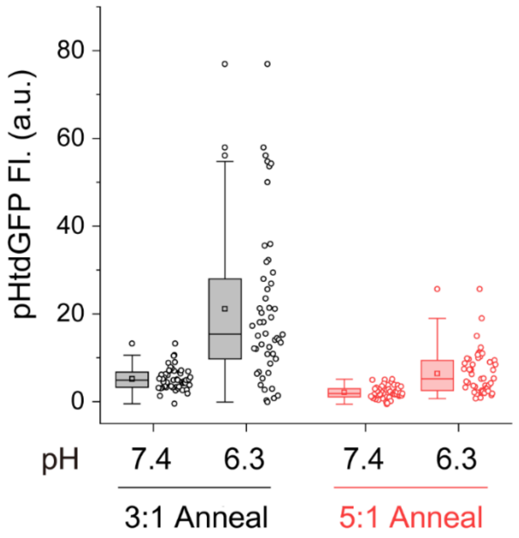


**Supplementary Fig. 18.** Effect of the pH-responsive ssDNA-to-trigger ratio on background and acid-activated pHtdGFP expression. Synthetic cells containing pH-responsive ssDNA and trigger strand at a 3:1 or 5:1 ratio were incubated under pH 7.4 or pH 6.3 conditions, and pHtdGFP fluorescence intensity was quantified for individual vesicles. Increasing the ratio from 3:1 to 5:1 reduced background expression at pH 7.4 but also decreased expression at pH 6.3, indicating a trade-off between leakage suppression and activated protein output. Individual vesicles are shown as data points ($n$ = 50 individual vesicles). Box plots indicate the median and interquartile range, squares indicate the mean, and whiskers extend to 1.5 times the interquartile range.

### Supplementary Tables

| Name | Sequence |
|---|---|
| pH-responsive ssDNA #1 | 5'- TTC TCT TCT CGT TTG CTC TTC TCT TGT GTG GGA TTG TCT AAG AGA AGA G |
| pH-responsive ssDNA #2 | 5'- TTC TCT TCT CGT TTG CTC TTC TCT TGT GTG GTA TTG TCC AAG AGA AGA G |
| pH-responsive ssDNA #3 | 5'- TTC TCT TCT CGT TTG CTC TTC TCT TGT GTG GTA TTG TTC AAG AGA AGA G |
| Trigger ssDNA #1 | 5'- TAT GCA AAC AAG ACA ATC CCA CAC AAG ATA AGA TTT TTT TTT T |
| Trigger ssDNA #2 | 5'- AGA CAA TCC CAC ACA ATT TTT TTT TT |
| Trigger ssDNA #3 | 5'- TAT GCA AAC AAG ACA ATA CCA CAC AAT TTT TTT TTT |
| Toehold switch RNA #1 | 5'- GG GUU GUG UGG GAU UGU CUU GUU UGC AUA CAG AAA CAG AGG AGA UAU GCA AUG AAG ACA AUA AAC CUG AAA GGA GCG CAA AAG AUG GUC UUC |
| Toehold switch RNA #2 | 5'- GG GUU GUG UGG UAU UGU CUU GUU UGC AUA CAG AAA CAG AGG AGA UAU GCA AUG AAG ACA AUA AAC CUG AAA GGA GCG CAA AAG AUG GUC UUC |

**Table S1. Sequences of the pH-responsive ssDNA, trigger ssDNA, and toehold switch RNA.** Toehold switch RNA sequences are shown up to 31 nucleotides following the end of the stem domain, including a 22-nt linker and the first 9 nucleotides of the NanoLuc (NLuc) coding region. Note that pH-responsive ssDNA #2 differs from #1 by a single base substitution (GGGATT → GGTATT). This modification was introduced with the intention of further reducing the hybridization affinity between the pH-responsive ssDNA and the trigger ssDNA. Hairpin-forming stem domains are underlined in both the pH-responsive ssDNA and toehold switch sequences.

| Annealing length | pH-responsive ssDNA | Trigger ssDNA | Toehold switch RNA |
|---|---|---|---|
| 18-nt | pH-responsive ssDNA #1 | Trigger ssDNA #1 | Toehold switch RNA #1 |
| 16-nt | pH-responsive ssDNA #1 | Trigger ssDNA #2 | Not tested* |
| 15-nt | pH-responsive ssDNA #2 | Trigger ssDNA #3 | Toehold switch RNA #2 |
| 14-nt | pH-responsive ssDNA #3 | Trigger ssDNA #3 | Toehold switch RNA #2 |

**Table S2. Sequence pairs for different annealing lengths shown in Fig. 1c and Fig. 2b.** Each sequence is shown in *Supplementary Table S1*. * Trigger ssDNA #2 is only for annealing experiments and not meant for toehold switch activation.

### Supplementary Video

**Supplementary Video 1. 3D visualization of Cy5-loaded synthetic cells embedded in alginate hydrogel.** Rotating 3D projection reconstructed from confocal z-stack images shows the spatial distribution of Cy5-containing vesicles throughout the hydrogel matrix. Scale bar, 10 μm.

### Supplementary Notes

#### 1. DNA sequences (plasmid and linear)

All the DNA sequences here are shown with the final toehold switch design (toehold switch #2). See *Supplementary Table S1* for the toehold switch #1 sequence. The toehold switch sequence is highlighted in red, pHtdGFP/sfGFP in green, NLuc in blue, and CPP in purple.

#### 1.1 Plasmid DNA sequences

##### 1.1.1 T7-toehold switch-pHtdGFP

CTCCGCTATCGCTACGTGACTGGGTCATGGCTGCGCCCCGACACCCGCCAACACCCGCTGACG
CGCCCTGACGGGCTTGTCTGCTCCCGGCATCCGCTTACAGACAAGCTGTGACCGTCTCCGGGA
GCTGCATGTGTCAGAGGTTTTCACCGTCATCACCGAAACGCGCGAGGCAGCTGCGGTAAAGCTC
ATCAGCGTGGTCGTGAAGCGATTCACAGATGTCTGCCTGTTCATCCGCGTCCAGCTCGTTGAGTT
TCTCCAGAAGCGTTAATGTCTGGCTTCTGATAAAGCGGGCCATGTTAAGGGCGGTTTTTTCCTGTT
TGGTCACTGATGCCTCCGTGTAAGGGGGATTTCTGTTCATGGGGGTAATGATACCGATGAAACGA
GAGAGGATGCTCACGATACGGGTTACTGATGATGAACATGCCCGGTTACTGGAACGTTGTGAGGG
TAAACAACTGGCGGTATGGATGCGGCGGGACCAGAGAAAAATCACTCAGGGTCAATGCCAGCGC
TTCGTTAATACAGATGTAGGTGTTCCACAGGGTAGCCAGCAGCATCCTGCGATGCAGATCCGGAA
CATAATGGTGCAGGGCGCTGACTTCCGCGTTTCCAGACTTTACGAAACACGGAAACCGAAGACC
ATTCATGTTGTTGCTCAGGTCGCAGACGTTTTGCAGCAGCAGTCGCTTCACGTTCGCTCGCGTAT
CGGTGATTCATTCTGCTAACCAGTAAGGCAACCCCGCCAGCCTAGCCGGGTCCTCAACGACAGG
AGCACGATCATGCGCACCCGTGGGGCCGCCATGCCGGCGATAATGGCCTGCTTCTCGCCGAAAC
GTTTGGTGGCGGGACCAGTGACGAAGGCTTGAGCGAGGGCGTGCAAGATTCCGAATACCGCAA
GCGACAGGCCGATCATCGTCGCGCTCCAGCGAAAGCGGTCCTCGCCGAAAATGACCCAGAGCG
CTGCCGGCACCTGTCCTACGAGTTGCATGATAAAGAAGACAGTCATAAGTGCGGCGACGATAGTC
ATGCCCCGCGCCCACCGGAAGGAGCTGACTGGGTTGAAGGCTCTCAAGGGCATCGGTCGAGAT
CCCGGTGCCTAATGAGTGAGCTAACTTACATTAATTGCGTTGCGCTCACTGCCCGCTTTCCAGTC
GGGAAACCTGTCGTGCCAGCTGCATTAATGAATCGGCCAACGCGCGGGGAGAGGCGGTTTGCG
TATTGGGCGCCAGGGTGGTTTTTCTTTTCACCAGTGAGACGGGCAACAGCTGATTGCCCTTCACC
GCCTGGCCCTGAGAGAGTTGCAGCAAGCGGTCCACGCTGGTTTGCCCCAGCAGGCGAAAATCC
TGTTTGATGGTGGTTAACGGCGGGATATAACATGAGCTGTCTTCGGTATCGTCGTATCCCACTACC
GAGATATCCGCACCAACGCGCAGCCCGGACTCGGTAATGGCGCGCATTGCGCCCAGCGCCATCT
GATCGTTGGCAACCAGCATCGCAGTGGGAACGATGCCCTCATTCAGCATTTGCATGGTTTGTTGA
AAACCGGACATGGCACTCCAGTCGCCTTCCCGTTCCGCTATCGGCTGAATTTGATTGCGAGTGAG
ATATTTATGCCAGCCAGCCAGACGCAGACGCGCCGAGACAGAACTTAATGGGCCCGCTAACAGC
GCGATTTGCTGGTGACCCAATGCGACCAGATGCTCCACGCCCAGTCGCGTACCGTCTTCATGGG
AGAAAATAATACTGTTGATGGGTGTCTGGTCAGAGACATCAAGAAATAACGCCGGAACATTAGTGC
AGGCAGCTTCCACAGCAATGGCATCCTGGTCATCCAGCGGATAGTTAATGATCAGCCCACTGACG
CGTTGCGCGAGAAGATTGTGCACCGCCGCTTTACAGGCTTCGACGCCGCTTCGTTCTACCATCG
ACACCACCACGCTGGCACCCAGTTGATCGGCGCGAGATTTAATCGCCGCGACAATTTGCGACGG
CGCGTGCAGGGCCAGACTGGAGGTGGCAACGCCAATCAGCAACGACTGTTTGCCCGCCAGTTG
TTGTGCCACGCGGTTGGGAATGTAATTCAGCTCCGCCATCGCCGCTTCCACTTTTTCCCGCGTTT
TCGCAGAAACGTGGCTGGCCTGGTTCACCACGCGGGAAACGGTCTGATAAGAGACACCGGCATA
CTCTGCGACATCGTATAACGTTACTGGTTTCACATTCACCACCCTGAATTGACTCTCTTCCGGGCG
CTATCATGCCATACCGCGAAAGGTTTTGCGCCATTCGATGGTGTCCGGGATCTCGACGCTCTCCC
TTATGCGACTCCTGCATTAGGAAGCAGCCCAGTAGTAGGTTGAGGCCGTTGAGCACCGCCGCCG
CAAGGAATGGTGCATGCAAGGAGATGGCGCCCAACAGTCCCCCGGCCACGGGGCCTGCCACCA
TACCCACGCCGAAACAAGCGCTCATGAGCCCGAAGTGGCGAGCCCGATCTTCCCCATCGGTGAT
GTCGGCGATATAGGCGCCAGCAACCGCACCTGTGGCGCCGGTGATGCCGGCCACGATGCGTCC
GGCGTAGAGGATCGAGATCTCGATCCCGCGAAATTAATACGACTCACTATAGGGTTGTGTGGTATT

GTCTTGTTTGCATACAGAAACAGAGGAGATATGCAATGAAGACAATAAACCTGAAAGGAGCGCAAA
AGatgcgtaaaggcgaagagctgttcacgggagtggtgccgattttagttgaactggatggggatgttaacggtcacaagtttagcgtgcgtg
gtgagggcgagggcgatgcgacgaatggtaaactgaccctgaaattcatctgtaccacaggcaaattgccagttccgtggccaacgctagtg
acgaccctgacttatggcgtccagtgcttttctaggtaccccgatcatatgaagcgccacgactttttaaaagcgctatgcccgaaggttacgtg
caagagcgcaccatcagcttcaaagatgacggcacgtacaaaactcgcgccgaagttaagttcgaaggtgataccctggttaatcgtatcga
gttgaagggtatcgacttcaaagaggatgggaatatcttgggccataaattggaatacaactttaactcacactatgtgtatattaccgcggataa
gcagaaaaacggtattaaggccaacttcaaaatccggcacaatgtagaagatggctcagttcagctcgcggatcattaccaacaaaacacc
ccgattggggatggtccggttcttctcccggataatcattacctttcgacccactctgttctcagcaaggacccgaacgaaaaacgcgatcacat
ggtgctgcttgaattcgtgactgcggctggaattactcatgggcatggaaccggcagcacgggctcggggtcatctggcacagcgagctcgg
aagataacaacatggcactgtttacaggcgtggtccctattctcgtagaattagacggtgacgtaaatggccacaaattcagtgtacgtggtga
gggcgaaggcgacgcaacaaacggcaaactgacgctgaaatttatatgcacgaccggcaagctgcccgttccatggccgaccttagtgac
caccttaacctacggtgtgcaatgttttagccgttatccggaccacatgaagcgccatgatttctttaaaagtgccatgccggagggatatgtcca
ggaacgaactatttcctttaaagacgatggcacttataaaacccgcgcggaagtaaaatttgagggtgacacgttggtcaatcgtattgaactg
aaaggcattgattttaaagaagatggcaacattttaggacacaaactagagtacaattttaattcgcattatgtctatatcaccgccgacaaaca
gaaaaatgggattaaagcgaatttcaaaattcgtcataacgtggaagatggtagtgtccagctggcagaccattatcagcagaacaccccga
ttggcgacggccctgtacttctgcctgataatcactatctgagcacgcattcagtgctgtccaaagatccgaacgaaaaacgcgatcatatggtc
ctgctggaatttgtgaccgcagccggaatcacacatggtatggatgaattatacaagcaccaccaccaccaccacTGAGATCCGGCT
GCTAACAAAGCCCGAAAGGAAGCTGAGTTGGCTGCTGCCACCGCTGAGCAATAACTAGCATAACC
CCTTGGGGCCTCTAAACGGGTCTTGAGGGGTTTTTTCTGAAAGGAGGAACTATATCCGGATTGGC
GAATGGGACGCGCCCTGTAGCGGCGCATTAAGCGCGGCGGGTGTGGTGGTTACGCGCAGCGTG
ACCGCTACACTTGCCAGCGCCCTAGCGCCCGCTCCTTTCGCTTTCTTCCCTTCCTTTCTCGCCAC
GTTCGCCGGCTTTCCCCGTCAAGCTCTAAATCGGGGGCTCCCTTTAGGGTTCCGATTTAGTGCTT
TACGGCACCTCGACCCCAAAAAACTTGATTAGGGTGATGGTTCACGTAGTGGGCCATCGCCCTGA
TAGACGGTTTTTCGCCCTTTGACGTTGGAGTCCACGTTCTTTAATAGTGGACTCTTGTTCCAAACT
GGAACAACACTCAACCCTATCTCGGTCTATTCTTTTGATTTATAAGGGATTTTGCCGATTTCGGCCT
ATTGGTTAAAAAATGAGCTGATTTAACAAAAATTTAACGCGAATTTTAACAAAATATTAACGTTTACAA
TTTCAGGTGGCACTTTTCGGGGAAATGTGCGCGGAACCCCTATTTGTTTATTTTTCTAAATACATTC
AAATATGTATCCGCTCATGAATTAATTCTTAGAAAAACTCATCGAGCATCAAATGAAACTGCAATTTA
TTCATATCAGGATTATCAATACCATATTTTTGAAAAAGCCGTTTCTGTAATGAAGGAGAAAACTCACC
GAGGCAGTTCCATAGGATGGCAAGATCCTGGTATCGGTCTGCGATTCCGACTCGTCCAACATCAA
TACAACCTATTAATTTCCCCTCGTCAAAAATAAGGTTATCAAGTGAGAAATCACCATGAGTGACGAC
TGAATCCGGTGAGAATGGCAAAAGTTTATGCATTTCTTTCCAGACTTGTTCAACAGGCCAGCCATT
ACGCTCGTCATCAAAATCACTCGCATCAACCAAACCGTTATTCATTCGTGATTGCGCCTGAGCGAG
ACGAAATACGCGATCGCTGTTAAAAGGACAATTACAAACAGGAATCGAATGCAACCGGCGCAGGA
ACACTGCCAGCGCATCAACAATATTTTCACCTGAATCAGGATATTCTTCTAATACCTGGAATGCTGT
TTTCCCGGGGATCGCAGTGGTGAGTAACCATGCATCATCAGGAGTACGGATAAAATGCTTGATGG
TCGGAAGAGGCATAAATTCCGTCAGCCAGTTTAGTCTGACCATCTCATCTGTAACATCATTGGCAA
CGCTACCTTTGCCATGTTTCAGAAACAACTCTGGCGCATCGGGCTTCCCATACAATCGATAGATTG
TCGCACCTGATTGCCCGACATTATCGCGAGCCCATTTATACCCATATAAATCAGCATCCATGTTGGA
ATTTAATCGCGGCCTAGAGCAAGACGTTTCCCGTTGAATATGGCTCATAACACCCCTTGTATTACTG
TTTATGTAAGCAGACAGTTTTATTGTTCATGACCAAAATCCCTTAACGTGAGTTTTCGTTCCACTGA
GCGTCAGACCCCGTAGAAAAGATCAAAGGATCTTCTTGAGATCCTTTTTTTCTGCGCGTAATCTGC
TGCTTGCAAACAAAAAAACCACCGCTACCAGCGGTGGTTTGTTTGCCGGATCAAGAGCTACCAAC
TCTTTTTCCGAAGGTAACTGGCTTCAGCAGAGCGCAGATACCAAATACTGTCCTTCTAGTGTAGCC
GTAGTTAGGCCACCACTTCAAGAACTCTGTAGCACCGCCTACATACCTCGCTCTGCTAATCCTGTT
ACCAGTGGCTGCTGCCAGTGGCGATAAGTCGTGTCTTACCGGGTTGGACTCAAGACGATAGTTAC
CGGATAAGGCGCAGCGGTCGGGCTGAACGGGGGGTTCGTGCACACAGCCCAGCTTGGAGCGA
ACGACCTACACCGAACTGAGATACCTACAGCGTGAGCTATGAGAAAGCGCCACGCTTCCCGAAG
GGAGAAAGGCGGACAGGTATCCGGTAAGCGGCAGGGTCGGAACAGGAGAGCGCACGAGGGAG
CTTCCAGGGGGAAACGCCTGGTATCTTTATAGTCCTGTCGGGTTTCGCCACCTCTGACTTGAGCG
TCGATTTTTGTGATGCTCGTCAGGGGGGCGGAGCCTATGGAAAAACGCCAGCAACGCGGCCTTT
TTACGGTTCCTGGCCTTTTGCTGGCCTTTTGCTCACATGTTCTTTCCTGCGTTATCCCCTGATTCT
GTGGATAACCGTATTACCGCCTTTGAGTGAGCTGATACCGCTCGCCGCAGCCGAACGACCGAGC
GCAGCGAGTCAGTGAGCGAGGAAGCGGAAGAGCGCCTGATGCGGTATTTTCTCCTTACGCATCT

GTGCGGTATTTCACACCGCATATATGGTGCACTCTCAGTACAATCTGCTCTGATGCCGCATAGTTAA
GCCAGTATACA

#### 1.1.2 T7-toehold switch-NLuc

CTCCGCTATCGCTACGTGACTGGGTCATGGCTGCGCCCCGACACCCGCCAACACCCGCTGACG
CGCCCTGACGGGCTTGTCTGCTCCCGGCATCCGCTTACAGACAAGCTGTGACCGTCTCCGGGA
GCTGCATGTGTCAGAGGTTTTCACCGTCATCACCGAAACGCGCGAGGCAGCTGCGGTAAAGCTC
ATCAGCGTGGTCGTGAAGCGATTCACAGATGTCTGCCTGTTCATCCGCGTCCAGCTCGTTGAGTT
TCTCCAGAAGCGTTAATGTCTGGCTTCTGATAAAGCGGGCCATGTTAAGGGCGGTTTTTTCCTGTT
TGGTCACTGATGCCTCCGTGTAAGGGGGATTTCTGTTCATGGGGGTAATGATACCGATGAAACGA
GAGAGGATGCTCACGATACGGGTTACTGATGATGAACATGCCCGGTTACTGGAACGTTGTGAGGG
TAAACAACTGGCGGTATGGATGCGGCGGGACCAGAGAAAAATCACTCAGGGTCAATGCCAGCGC
TTCGTTAATACAGATGTAGGTGTTCCACAGGGTAGCCAGCAGCATCCTGCGATGCAGATCCGGAA
CATAATGGTGCAGGGCGCTGACTTCCGCGTTTCCAGACTTTACGAAACACGGAAACCGAAGACC
ATTCATGTTGTTGCTCAGGTCGCAGACGTTTTGCAGCAGCAGTCGCTTCACGTTCGCTCGCGTAT
CGGTGATTCATTCTGCTAACCAGTAAGGCAACCCCGCCAGCCTAGCCGGGTCCTCAACGACAGG
AGCACGATCATGCGCACCCGTGGGGCCGCCATGCCGGCGATAATGGCCTGCTTCTCGCCGAAAC
GTTTGGTGGCGGGACCAGTGACGAAGGCTTGAGCGAGGGCGTGCAAGATTCCGAATACCGCAA
GCGACAGGCCGATCATCGTCGCGCTCCAGCGAAAGCGGTCCTCGCCGAAAATGACCCAGAGCG
CTGCCGGCACCTGTCCTACGAGTTGCATGATAAAGAAGACAGTCATAAGTGCGGCGACGATAGTC
ATGCCCCGCGCCCACCGGAAGGAGCTGACTGGGTTGAAGGCTCTCAAGGGCATCGGTCGAGAT
CCCGGTGCCTAATGAGTGAGCTAACTTACATTAATTGCGTTGCGCTCACTGCCCGCTTTCCAGTC
GGGAAACCTGTCGTGCCAGCTGCATTAATGAATCGGCCAACGCGCGGGGAGAGGCGGTTTGCG
TATTGGGCGCCAGGGTGGTTTTTCTTTTCACCAGTGAGACGGGCAACAGCTGATTGCCCTTCACC
GCCTGGCCCTGAGAGAGTTGCAGCAAGCGGTCCACGCTGGTTTGCCCCAGCAGGCGAAAATCC
TGTTTGATGGTGGTTAACGGCGGGATATAACATGAGCTGTCTTCGGTATCGTCGTATCCCACTACC
GAGATATCCGCACCAACGCGCAGCCCGGACTCGGTAATGGCGCGCATTGCGCCCAGCGCCATCT
GATCGTTGGCAACCAGCATCGCAGTGGGAACGATGCCCTCATTCAGCATTTGCATGGTTTGTTGA
AAACCGGACATGGCACTCCAGTCGCCTTCCCGTTCCGCTATCGGCTGAATTTGATTGCGAGTGAG
ATATTTATGCCAGCCAGCCAGACGCAGACGCGCCGAGACAGAACTTAATGGGCCCGCTAACAGC
GCGATTTGCTGGTGACCCAATGCGACCAGATGCTCCACGCCCAGTCGCGTACCGTCTTCATGGG
AGAAAATAATACTGTTGATGGGTGTCTGGTCAGAGACATCAAGAAATAACGCCGGAACATTAGTGC
AGGCAGCTTCCACAGCAATGGCATCCTGGTCATCCAGCGGATAGTTAATGATCAGCCCACTGACG
CGTTGCGCGAGAAGATTGTGCACCGCCGCTTTACAGGCTTCGACGCCGCTTCGTTCTACCATCG
ACACCACCACGCTGGCACCCAGTTGATCGGCGCGAGATTTAATCGCCGCGACAATTTGCGACGG
CGCGTGCAGGGCCAGACTGGAGGTGGCAACGCCAATCAGCAACGACTGTTTGCCCGCCAGTTG
TTGTGCCACGCGGTTGGGAATGTAATTCAGCTCCGCCATCGCCGCTTCCACTTTTTCCCGCGTTT
TCGCAGAAACGTGGCTGGCCTGGTTCACCACGCGGGAAACGGTCTGATAAGAGACACCGGCATA
CTCTGCGACATCGTATAACGTTACTGGTTTCACATTCACCACCCTGAATTGACTCTCTTCCGGGCG
CTATCATGCCATACCGCGAAAGGTTTTGCGCCATTCGATGGTGTCCGGGATCTCGACGCTCTCCC
TTATGCGACTCCTGCATTAGGAAGCAGCCCAGTAGTAGGTTGAGGCCGTTGAGCACCGCCGCCG
CAAGGAATGGTGCATGCAAGGAGATGGCGCCCAACAGTCCCCCGGCCACGGGGCCTGCCACCA
TACCCACGCCGAAACAAGCGCTCATGAGCCCGAAGTGGCGAGCCCGATCTTCCCCATCGGTGAT
GTCGGCGATATAGGCGCCAGCAACCGCACCTGTGGCGCCGGTGATGCCGGCCACGATGCGTCC
GGCGTAGAGGATCGAGATCTCGATCCCGCGAAATTAATACGACTCACTATAGGGTTGTGTGGTATT
GTCTTGTTTGCATACAGAAACAGAGGAGATATGCAATGAAGACAATAAACCTGAAAGGAGCGCAAA
AGatggtcttcacactcgaagatttcgttggggactgggaacagacagccgcctacaacctggaccaagtccttgaacagggaggtgtgtcc
agtttgctgcagaatctcgccgtgtccgtaactccgatccaaaggattgtccggagcggtgaaaatgccctgaagatcgacatccatgtcatcat
cccgtatgaaggtctgagcgccgaccaaatggcccagatcgaagaggtgtttaaggtggtgtaccctgtggatgatcatcactttaaggtgatc
ctgccctatggcacactggtaatcgacggggttacgccgaacatgctgaactatttcggacggccgtatgaaggcatcgccgtgttcgacggc
aaaaagatcactgtaacagggaccctgtggaacggcaacaaaattatcgacgagcgcctgatcacccccgacggctccatgctgttccgag
taaccatcaacagcgtgaccggctaccggctgttcgaggagattctgagcggttctcaccaccaccaccaccacTGAGATCCGGCT

GCTAACAAAGCCCGAAAGGAAGCTGAGTTGGCTGCTGCCACCGCTGAGCAATAACTAGCATAACC
CCTTGGGGCCTCTAAACGGGTCTTGAGGGGTTTTTTCTGAAAGGAGGAACTATATCCGGATTGGC
GAATGGGACGCGCCCTGTAGCGGCGCATTAAGCGCGGCGGGTGTGGTGGTTACGCGCAGCGTG
ACCGCTACACTTGCCAGCGCCCTAGCGCCCGCTCCTTTCGCTTTCTTCCCTTCCTTTCTCGCCAC
GTTCGCCGGCTTTCCCCGTCAAGCTCTAAATCGGGGGCTCCCTTTAGGGTTCCGATTTAGTGCTT
TACGGCACCTCGACCCCAAAAAACTTGATTAGGGTGATGGTTCACGTAGTGGGCCATCGCCCTGA
TAGACGGTTTTTCGCCCTTTGACGTTGGAGTCCACGTTCTTTAATAGTGGACTCTTGTTCCAAACT
GGAACAACACTCAACCCTATCTCGGTCTATTCTTTTGATTTATAAGGGATTTTGCCGATTTCGGCCT
ATTGGTTAAAAAATGAGCTGATTTAACAAAAATTTAACGCGAATTTTAACAAAATATTAACGTTTACAA
TTTCAGGTGGCACTTTTCGGGGAAATGTGCGCGGAACCCCTATTTGTTTATTTTTCTAAATACATTC
AAATATGTATCCGCTCATGAATTAATTCTTAGAAAAACTCATCGAGCATCAAATGAAACTGCAATTTA
TTCATATCAGGATTATCAATACCATATTTTTGAAAAAGCCGTTTCTGTAATGAAGGAGAAAACTCACC
GAGGCAGTTCCATAGGATGGCAAGATCCTGGTATCGGTCTGCGATTCCGACTCGTCCAACATCAA
TACAACCTATTAATTTCCCCTCGTCAAAAATAAGGTTATCAAGTGAGAAATCACCATGAGTGACGAC
TGAATCCGGTGAGAATGGCAAAAGTTTATGCATTTCTTTCCAGACTTGTTCAACAGGCCAGCCATT
ACGCTCGTCATCAAAATCACTCGCATCAACCAAACCGTTATTCATTCGTGATTGCGCCTGAGCGAG
ACGAAATACGCGATCGCTGTTAAAAGGACAATTACAAACAGGAATCGAATGCAACCGGCGCAGGA
ACACTGCCAGCGCATCAACAATATTTTCACCTGAATCAGGATATTCTTCTAATACCTGGAATGCTGT
TTTCCCGGGGATCGCAGTGGTGAGTAACCATGCATCATCAGGAGTACGGATAAAATGCTTGATGG
TCGGAAGAGGCATAAATTCCGTCAGCCAGTTTAGTCTGACCATCTCATCTGTAACATCATTGGCAA
CGCTACCTTTGCCATGTTTCAGAAACAACTCTGGCGCATCGGGCTTCCCATACAATCGATAGATTG
TCGCACCTGATTGCCCGACATTATCGCGAGCCCATTTATACCCATATAAATCAGCATCCATGTTGGA
ATTTAATCGCGGCCTAGAGCAAGACGTTTCCCGTTGAATATGGCTCATAACACCCCTTGTATTACTG
TTTATGTAAGCAGACAGTTTTATTGTTCATGACCAAAATCCCTTAACGTGAGTTTTCGTTCCACTGA
GCGTCAGACCCCGTAGAAAAGATCAAAGGATCTTCTTGAGATCCTTTTTTTCTGCGCGTAATCTGC
TGCTTGCAAACAAAAAAACCACCGCTACCAGCGGTGGTTTGTTTGCCGGATCAAGAGCTACCAAC
TCTTTTTCCGAAGGTAACTGGCTTCAGCAGAGCGCAGATACCAAATACTGTCCTTCTAGTGTAGCC
GTAGTTAGGCCACCACTTCAAGAACTCTGTAGCACCGCCTACATACCTCGCTCTGCTAATCCTGTT
ACCAGTGGCTGCTGCCAGTGGCGATAAGTCGTGTCTTACCGGGTTGGACTCAAGACGATAGTTAC
CGGATAAGGCGCAGCGGTCGGGCTGAACGGGGGGTTCGTGCACACAGCCCAGCTTGGAGCGA
ACGACCTACACCGAACTGAGATACCTACAGCGTGAGCTATGAGAAAGCGCCACGCTTCCCGAAG
GGAGAAAGGCGGACAGGTATCCGGTAAGCGGCAGGGTCGGAACAGGAGAGCGCACGAGGGAG
CTTCCAGGGGGAAACGCCTGGTATCTTTATAGTCCTGTCGGGTTTCGCCACCTCTGACTTGAGCG
TCGATTTTTGTGATGCTCGTCAGGGGGGCGGAGCCTATGGAAAAACGCCAGCAACGCGGCCTTT
TTACGGTTCCTGGCCTTTTGCTGGCCTTTTGCTCACATGTTCTTTCCTGCGTTATCCCCTGATTCT
GTGGATAACCGTATTACCGCCTTTGAGTGAGCTGATACCGCTCGCCGCAGCCGAACGACCGAGC
GCAGCGAGTCAGTGAGCGAGGAAGCGGAAGAGCGCCTGATGCGGTATTTTCTCCTTACGCATCT
GTGCGGTATTTCACACCGCATATATGGTGCACTCTCAGTACAATCTGCTCTGATGCCGCATAGTTAA
GCCAGTATACA

### 1.2 Linear DNA sequences

#### 1.2.1. T7-toehold switch-sfGFP-His

TCCGGCGTAGAGGATCGAGATCTCGATCCCGCGAAATTAATACGACTCACTATAGGGTTGTGTGGT
ATTGTCTTGTTTGCATACAGAAACAGAGGAGATATGCAATGAAGACAATAAACCTGAAAGGAGCGC
AAAAGatgcgtaaaggcgaagagctgttcactggtgtcgtccctattctggtggaactggatggtgatgtcaacggtcataagttttccgtgcgt
ggcgagggtgaaggtgacgcaactaatggtaaactgacgctgaagttcatctgtactactggtaaactgccggtaccttggccgactctggtaa
cgacgctgacttatggtgttcagtgctttgctcgttatccggaccatatgaagcagcatgacttcttcaagtccgccatgccggaaggctatgtgca
ggaacgcacgatttcctttaaggatgacggcacgtacaaaacgcgtgcggaagtgaaatttgaaggcgataccctggtaaaccgcattgagc
tgaaaggcattgactttaaagaagacggcaatatcctgggccataagctggaatacaattttaacagccacaatgtttacatcaccgccgataa
acaaaaaaatggcattaaagcgaattttaaaattcgccacaacgtggaggatggcagcgtgcagctggctgatcactaccagcaaaacact
ccaatcggtgatggtcctgttctgctgccagacaatcactatctgagcacgcaaagcgttctgtctaaagatccgaacgagaaacgcgatcata
tggttctgctggagttcgtaaccgcagcgggcatcacgcatggtatggatgaactatacaaacaccaccaccaccaccacTGAGATCC
GGCTGCTAACAAAGCCCGAAAGGAAGCTGAGTTGGCTGCTGCCACCGCTGAGCAATAACTAGCA
TAACCCCTTGGGGCCTCTAAACGGGTCTTGAGGGGTTTTTTCTGAAAGGAGGAACTATATCCGGA
TTGGCGAATGGGACG

#### 1.2.2. T7-toehold switch-CPP-linker-sfGFP-His

TCCGGCGTAGAGGATCGAGATCTCGATCCCGCGAAATTAATACGACTCACTATAGGGTTGTGTGGT
ATTGTCTTGTTTGCATACAGAAACAGAGGAGATATGCAATGAAGACAATAAACCTGAAAGGAGCGC
AAAAGatgcgtcaaataaagatttggtttcaaaatcgtcggatgaagtggaagaagggcggcggatcgggtggcggtagcatgcgtaaag
gcgaagagctgttcactggtgtcgtccctattctggtggaactggatggtgatgtcaacggtcataagttttccgtgcgtggcgagggtgaaggtg
acgcaactaatggtaaactgacgctgaagttcatctgtactactggtaaactgccggtaccttggccgactctggtaacgacgctgacttatggt
gttcagtgctttgctcgttatccggaccatatgaagcagcatgacttcttcaagtccgccatgccggaaggctatgtgcaggaacgcacgatttcc
tttaaggatgacggcacgtacaaaacgcgtgcggaagtgaaatttgaaggcgataccctggtaaaccgcattgagctgaaaggcattgacttt
aaagaagacggcaatatcctgggccataagctggaatacaattttaacagccacaatgtttacatcaccgccgataaacaaaaaaatggcat
taaagcgaattttaaaattcgccacaacgtggaggatggcagcgtgcagctggctgatcactaccagcaaaacactccaatcggtgatggtc
ctgttctgctgccagacaatcactatctgagcacgcaaagcgttctgtctaaagatccgaacgagaaacgcgatcatatggttctgctggagttc
gtaaccgcagcgggcatcacgcatggtatggatgaactatacaaacaccaccaccaccaccacTGAGATCCGGCTGCTAACA
AAGCCCGAAAGGAAGCTGAGTTGGCTGCTGCCACCGCTGAGCAATAACTAGCATAACCCCTTGG
GGCCTCTAAACGGGTCTTGAGGGGTTTTTTCTGAAAGGAGGAACTATATCCGGATTGGCGAATGG
GACG
